# Cold-Induced Hepatocyte-Derived Exosomes Activate Brown Adipose Thermogenesis via miR-293-5p-Mediated Transcriptional Reprogramming

**DOI:** 10.1101/2025.04.04.647213

**Authors:** Xiujuan Gao, Junqing Xu, Zengqiang Xu, Mengxin Jiang, Jiahao Zhu, Yang Geng, Shengjun Dong, Yanna Li, Zhengtong Zhou, Yingjiang Xu

**Author notes:** Correspondence, Binzhou Medical University, 346 Guanhai Rd, Yantai City, 264003, P.R.China. Institute of Medical Genomics, Biomedical Sciences College & Shandong Medicinal Biotechnology Centre, Medical Science and Technology Innovation Center, Shandong First Medical University & Shandong Academy of Medical Sciences, Jinan City, 271016, P.R.China. Binzhou Medical University Hospital, 661 Huanghe 2rd Rd, Binzhou City, 256603, P.R.China. These authors contributed equally (Xiujuan Gao, Junqing Xu, Zengqiang Xu).

## Abstract

The liver-adipose axis serves as a fundamental regulatory network governing systemic lipid homeostasis, with hepatic-derived signals orchestrating adipose tissue plasticity through multimodal mechanisms. Comprehensive elucidation of these bidirectional crosstalk mechanisms may reveal novel therapeutic strategies for metabolic disorders. Our investigation demonstrates that cold exposure induces hepatic secretion of exosomes that potentiate adipose thermogenic activation in both in vitro and in vivo models. This thermogenic potentiation is mechanistically linked to cold-induced upregulation of hepatocyte-derived exosomal miR-293-5p. Notably, pharmacological administration of miR-293-5p agomir significantly attenuates diet-induced obesity and associated metabolic dysregulation in murine models. Through mechanistic interrogation, we identified *Tet1* as a direct downstream target of miR- 293-5p, where ectopic *Tet1* expression disrupts brown adipose tissue (BAT) thermogenic programming independently of miR-293-5p modulation. Our findings establish cold-activated hepatocyte exosomes as endocrine signaling mediators containing thermogenic microRNA cargos, with miR-293-5p emerging as a regulatory nexus coordinating mitochondrial bioenergetics and systemic energy equilibrium through liver-adipose cross-talk.

## Background

Adipose tissue, a dynamic endocrine organ, orchestrates systemic energy homeostasis through its dual functional components. Brown adipose tissue (BAT) exhibits distinctive morphological features including dense vascular networks, extensive sympathetic innervation, and multilocular adipocytes replete with mitochondria-enriched cytoplasm. The thermogenic signature of BAT is defined by uncoupling protein 1 (UCP1)-mediated nonshivering thermogenesis, which dissipates chemical energy as heat through mitochondrial proton leakage [1–3]. In contrast, white adipose tissue (WAT) demonstrates remarkable plasticity through its capacity for "browning" - a cold/exercise/agonist-inducible process generating UCP1^+^ beige adipocytes that express canonical BAT markers (*Cidea, Cox7a, Prdm16, Pgc-1α*) and acquire thermogenic competence [4,5]. This adaptive thermogenic reprogramming, characterized by metabolic flexibility and inducibility, presents promising therapeutic potential for combating metabolic disorders.

The autonomic nervous system serves as the principal regulator of BAT activation, with cold-induced sympathetic stimulation triggering norepinephrine release and subsequent β3-adrenergic receptor signaling [6]. Beyond classical adrenergic pathways, emerging evidence highlights a complex regulatory network involving paracrine/endocrine factors and metabolite signaling that coordinates inter-tissue communication [7]. While extensive research has delineated transcriptional cascades downstream of β3-AR activation, the spatiotemporal integration of non-canonical signaling modalities remains incompletely characterized.

Extracellular vesicles (EVs), particularly exosomes, have emerged as critical mediators of interorgan crosstalk in metabolic regulation. These nano-scale carriers facilitate miRNA transfer between tissues, with their cargo profiles reflecting physiological status. Obesogenic conditions markedly alter plasma exosomal miRNA signatures (e.g., upregulated miR-122, miR-192, miR-27a-3p/b-3p), which can transfer metabolic dysfunction to recipient tissues [11]. Adipocyte-specific Dicer knockout models (ADicerKD) and human lipodystrophy cases demonstrate that adipose-derived exosomal miRNAs are indispensable for physiological tissue remodeling [12]. Notably, endogenous adipocyte miRNAs like miR-34a-5p and miR-155 differentially regulate brown/beige adipogenesis through targeting Fgfr1/Klb/Sirt1 and modulating proliferation-differentiation transitions, respectively [13,14]. The potential extra-adipose origins of these regulatory miRNAs via exosomal trafficking warrant further investigation.

As the central metabolic hub, the liver dynamically coordinates nutrient partitioning through intricate regulation of glucose and lipid fluxes [15,16]. Our study elucidates a previously unrecognized liver-BAT communication axis mediated by hepatocyte-derived exosomes (hepExos). Cold exposure induces hepatic secretion of exosomes enriched with miR-293-5p, which enhances BAT thermogenic capacity both in vitro and in vivo. Integrated miRNA sequencing and computational analyses identified Ten-eleven translocation methylcytosine dioxygenase 1 (Tet1) as a direct target of miR-293-5p. This discovery of the hepExo-miRNA-Tet1 regulatory axis provides novel mechanistic insights into interorgan metabolic communication and identifies potential therapeutic targets for metabolic syndrome.

## Results

### Cold-Activated Hepatocyte-Derived Exosomes Promote BAT Thermogenic Activation

To investigate the thermogenic regulatory potential of hepatic exosomes, 10- week-old C57BL/6 mice under standard diet were exposed to 4°C or room temperature (RT) for 3 days. Primary hepatocytes were isolated from liver tissues, and hepatocyte-derived exosomes (hepExos) were purified using an optimized protocol (Supp. Fig 1A). Quantification revealed significantly higher exosome yields from cold-exposed hepatocytes versus RT controls (Supp. Fig 1B). Transmission electron microscopy (TEM) confirmed the characteristic cup-shaped morphology of isolated vesicles (Supp. Fig 1C), while western blot (Wb) validated enrichment of exosomal markers CD9 and TSG101 (Supp. Fig 1D), collectively confirming successful hepExos isolation.

Ten-week-old male C57BL/6 mice maintained on a standard chow diet were subjected to 72-hour cold acclimation (4°C) or room temperature (25°C) exposure. Primary hepatocytes were isolated via collagenase perfusion digestion and cultured under serum-free conditions. HepExos were harvested from conditioned media using sequential ultracentrifugation and sucrose density gradient purification (Supp. Fig 1A). Quantitative analysis revealed a 1.8-fold increase in exosome yield from cold-exposed hepatocytes compared to controls (Supp. Fig 1B). Transmission electron microscopy confirmed the presence of 80-150 nm cup-shaped vesicles with double-membrane structures (Supp. Fig 1C). Immunoblotting demonstrated robust enrichment of canonical exosomal markers, including CD9 and TSG101, with negligible contamination from cellular organelle proteins (Supp. Fig 1D). This protocol establishes the methodology’s robustness for isolating intact hepExos with preserved structural integrity and molecular signatures.

As nanoscale mediators of intercellular communication, exosomes orchestrate metabolic coordination through transfer of bioactive cargo including miRNAs, proteins, and lipids [17,18]. To investigate the biodynamics of hepExos, PKH26- labeled particles (1×10^9^/mouse) were intravenously administered to 10-week-old C57BL/6J wild-type (WT) mice. Fluorescence microscopy at the 16-hour post-injection window revealed selective biodistribution, with hepatic and BAT tropism (Fig.1A). Notably, inguinal white adipose tissue (iWAT) exhibited undetectable signal intensity, demonstrating tissue-specific exosomal trafficking. This evidence conclusively establishes BAT as a privileged recipient of circulating hepExos.

**Figure 1.**
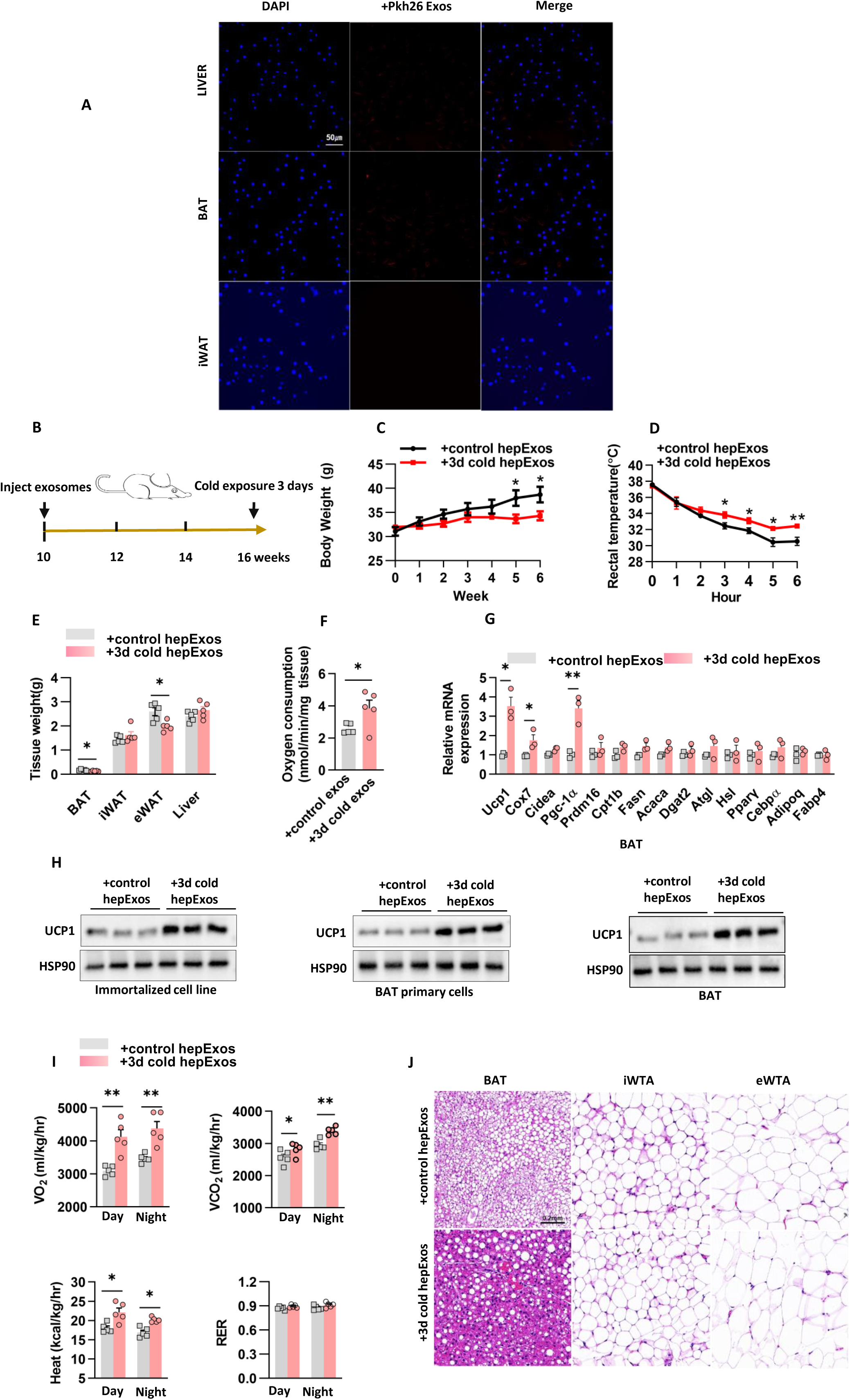
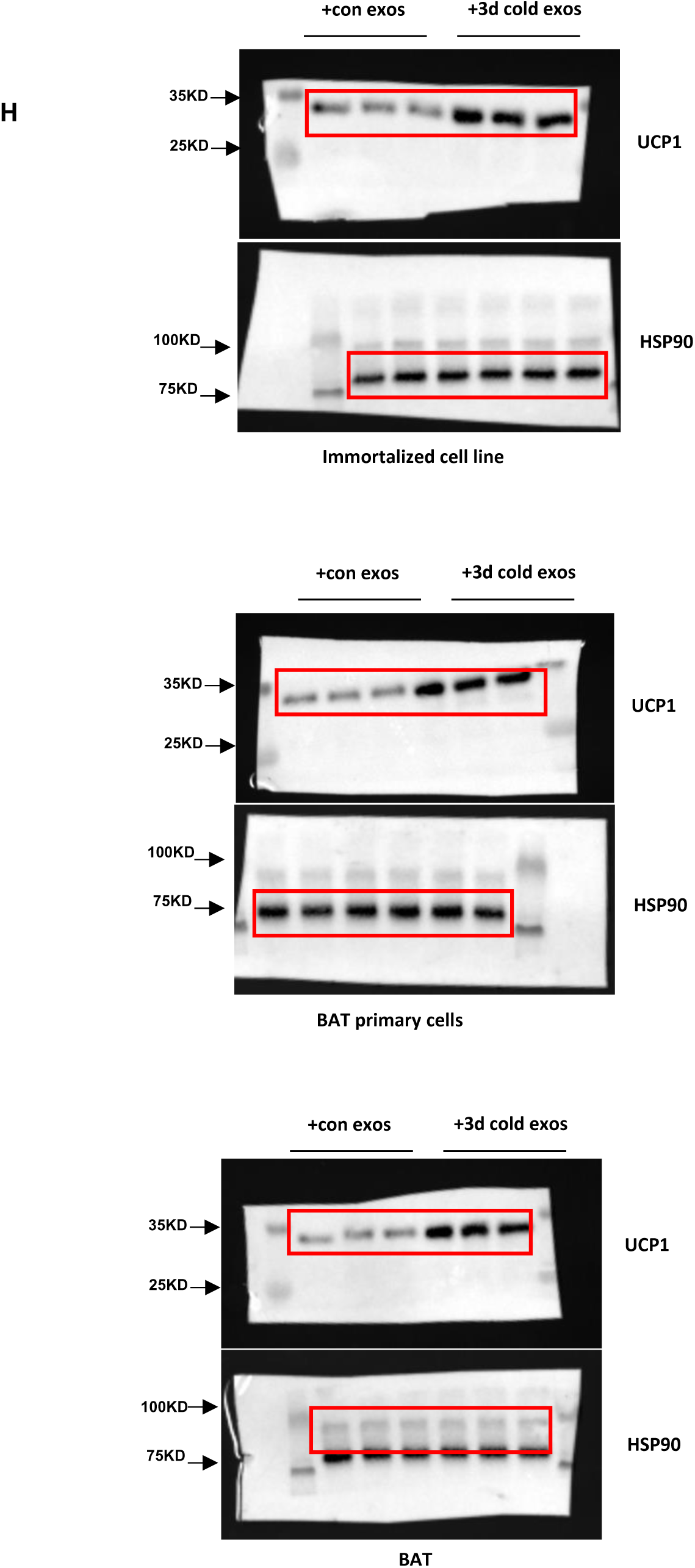
Cold-Activated Hepatocyte-Derived Exosomes Promote BAT Thermogenic Activation. A. HepExos could be transported to target BAT. Scale bar: 200μm. B. The in vivo protocol that used tail vein injection cold-exposed exosomes. C. Body weight of mice in (B). n=5 mice each group. D. Rectal temperature was measured when mice in (B) were challenged to 4°C. n=5 mice each group. E. Fat and liver mass weight in mice of (B). n=5 mice each group. F. Oxygen consumption rate measured by Clark electrodes for BAT as in (B). n=5 independent experiments. G. Gene expression analyzed by quantitative PCR in BAT as in (B). n=5 independent experiments. H. UCP1 protein levels were analyzed in immortalized brown adipocytes, BAT primary cells and BAT. n=3 independent experiments. I. C57BL/6J male mice were subjected to the indirect calorimetry analysis at basal state. n=5 mice each group. J. Hematoxylin and eosin staining of BAT of mice in (B). Scale bar: 0.2mm for BAT, iWAT and eWAT.

To investigate the potential involvement of hepExos in BAT) thermogenic regulation, we administered hepExos (5×10⁹ particles/mouse, twice weekly for 6 weeks) intravenously to 10-week-old WT mice, with exosomes from room temperature-adapted mice serving as controls (Fig. 1B). Remarkably, mice receiving cold-adapted hepExos (3-day 4°C exposure) demonstrated attenuated body weight gain (Fig. 1C) and enhanced cold tolerance (Fig. 1D), despite comparable food/water intake and serum triglyceride levels between groups (Supp. Figs. 2A-C). The metabolic improvements were accompanied by significant reductions in BAT mass and epididymal white adipose tissue (eWAT) weight (Fig. 1E), coupled with elevated BAT oxygen consumption rates (Fig. 1F). Molecular analysis revealed cold-adapted hepExos specifically upregulated thermogenic gene expression in BAT (*Ucp1, Cox7a, Pgc-1α*) without affecting iWAT (Fig. 1G, Supp. Fig. 2D). Consistently, UCP1 protein levels increased significantly in BAT (Fig. 1H), while remaining unchanged in iWAT (Supp. Fig. 2E). Whole-animal metabolic profiling through indirect calorimetry revealed that cold-adapted hepExos treatment elevated basal heat production , oxygen consumption, and CO₂ output (Fig. 1I). Histological analysis demonstrated BAT morphological remodeling with reduced lipid droplet size (Fig. 1J), whereas iWAT and eWAT adipocyte morphology remained unaltered.

Complementing these in vivo observations, we systematically evaluated the cell-autonomous effects of cold-adapted hepExos in vitro. Following standardized protocol, immortalized brown adipocytes (iBAs) and primary brown adipocytes were treated with cold-exposed hepExos (1×10⁸ particles per 0.5×10⁶ cells) in exosome-free differentiation medium for 48 hr. Remarkably, qPCR analysis revealed upregulation of core thermogenic effectors across all adipocyte models (Fig. 1H). Collectively, these results demonstrate that hepExos from cold-exposed mice enhance the thermogenic program in BAT.

### Hepatocyte exosomal miRNAs drive interorgan metabolic crosstalk

Building on established evidence of YBX1’s critical role in exosomal miRNA packaging [19,20], we employed liver-specific *Ybx1* knockdown to interrogate miRNA functionality in hepExos. C57BL/6 mice received tail vein injections of AAV9-shYbx1 or control AAV9-Ctrl (2×10¹¹ vector genomes in 200 μL PBS, with booster injections every 4 weeks). Following 10 weeks of standard chow diet and 3- day cold acclimation (4°C) (Supp. Fig. 3A), quantitative RT-PCR demonstrated approximately 70% reduction in hepatic Ybx1 mRNA (Supp. Fig. 3B). Crucially, *Ybx1* knocking down did not affect hepExos production (Supp. Fig. 3C), while miRNA quantification showed 80% depletion in hepExos, as evidenced by miR-122 levels - a hepatocyte-enriched miRNA serving as cargo biomarker (Supp. Fig. 3D).

Subsequent cold tolerance assessments in AAV9-YBX1 knockdown (KD) mice demonstrated significantly lower core body temperatures compared to AAV9-Ctrl KD mice (Supp. Fig. 3E). *Ybx1* deficiency resulted in a increase in body weight (Supp. Fig. 3F) and elevated serum triglyceride levels (Supp. Fig. 3G). Notably, BAT oxygen consumption rates decreased in KD mice (Supp. Fig. 3H). Transcriptional profiling revealed impaired thermogenesis in BAT, with *Ucp1, Cidea, Pgc-1α*, and *Prdm16* expression reduced by 30-52% (Supp. Fig. 3I). This molecular suppression correlated with a reduction in UCP1 protein abundance (Supp. Fig. 3J). Comprehensive metabolic analyses confirmed systemic impairments, showing also decrease in heat production, VO₂ , and VCO₂ (Supp. Fig. 3K). Histologically, BAT from KD mice exhibited 1-fold larger lipid droplets, while white adipose tissue morphology remained comparable between groups (Supp. Fig. 3L). Collectively, these findings establish YBX1 as a critical regulator of hepatic exosome-mediated thermogenic programming in BAT during cold adaptation.

Consistent with endogenous *Ybx1* deficiency, administration of AAV9-YBX1 KD hepExos profoundly disrupted BAT thermogenic programming. In our experimental paradigm (Fig. 2A), wild-type mice receiving 5×10^9^ KD hepExos via biweekly intravenous injections over six weeks showed comparable ad libitum consumption and serum triglycerides to controls (Supp. Figs. 4A-C). Strikingly, YBX1 KD hepExos-treated mice exhibited accelerated weight gain and impaired cold tolerance (Figs. 2B,C). This metabolic perturbation correlated with elevated ectopic fat deposition, particularly in epididymal WAT (eWAT) mass (Fig. 2D). Molecular profiling revealed systemic suppression of thermogenic machinery, evidenced reductions in BAT *Ucp1, Pgc-1α* and *Cidea* mRNA (Figs. 2E,F), paralleled decreased in UCP1 protein expression (Fig. 2G). Notably, these effects were BAT-specific, with iWAT morphology and gene expression remaining unaffected (Supp. Figs. 4D,E). Indirect calorimetry quantified the metabolic deficit, showing lower VO₂ and reduced heat production (Fig. 2H). Histologically, BAT from KD hepExos recipients displayed hypertrophic lipid dropletsand fewer UCP1^+^ multilocular adipocytes (Fig. 2I). These findings mechanistically establish miRNAs as indispensable effector molecules mediating hepExos’ regulation of adaptive thermogenesis.

**Figure 2.**
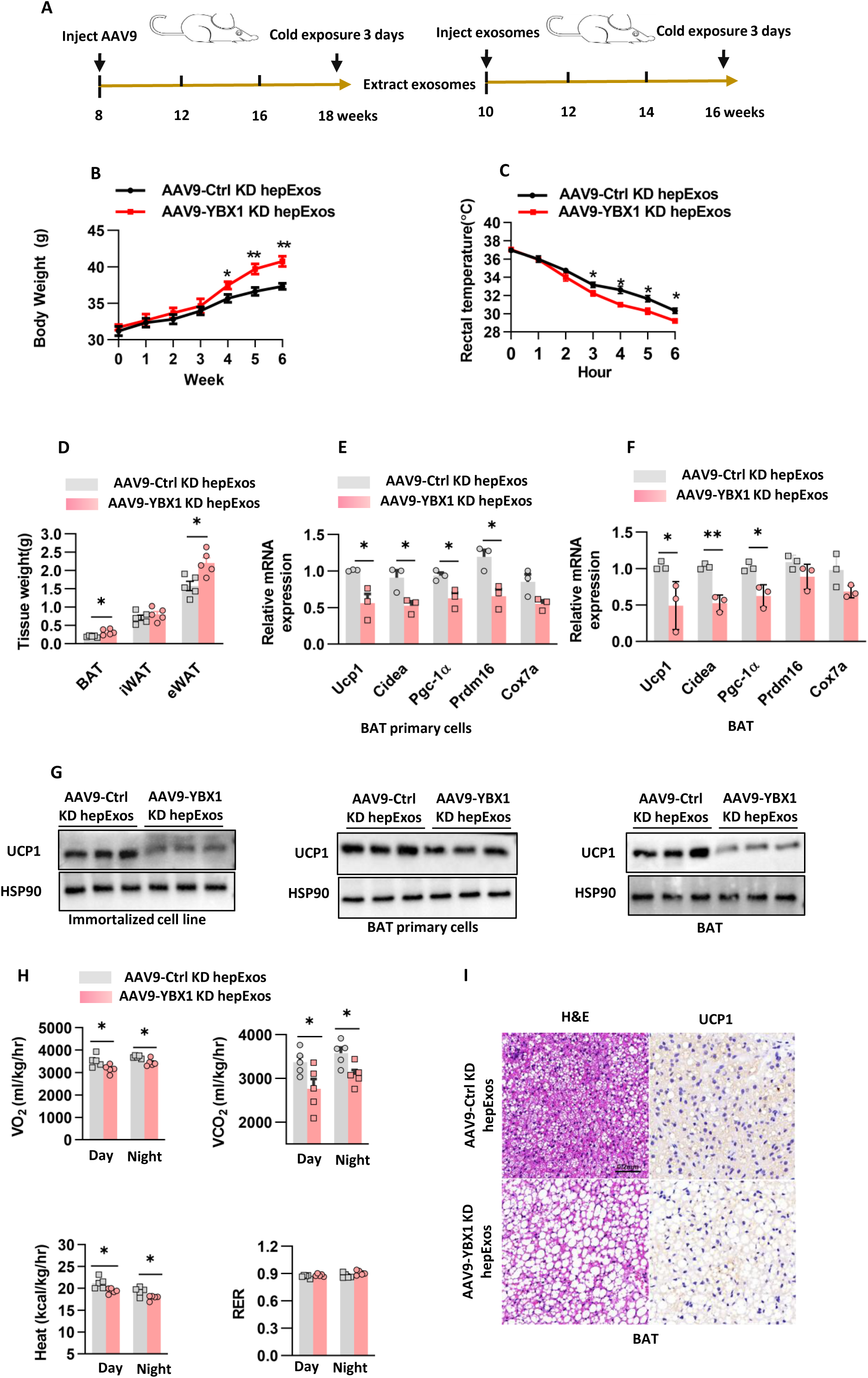
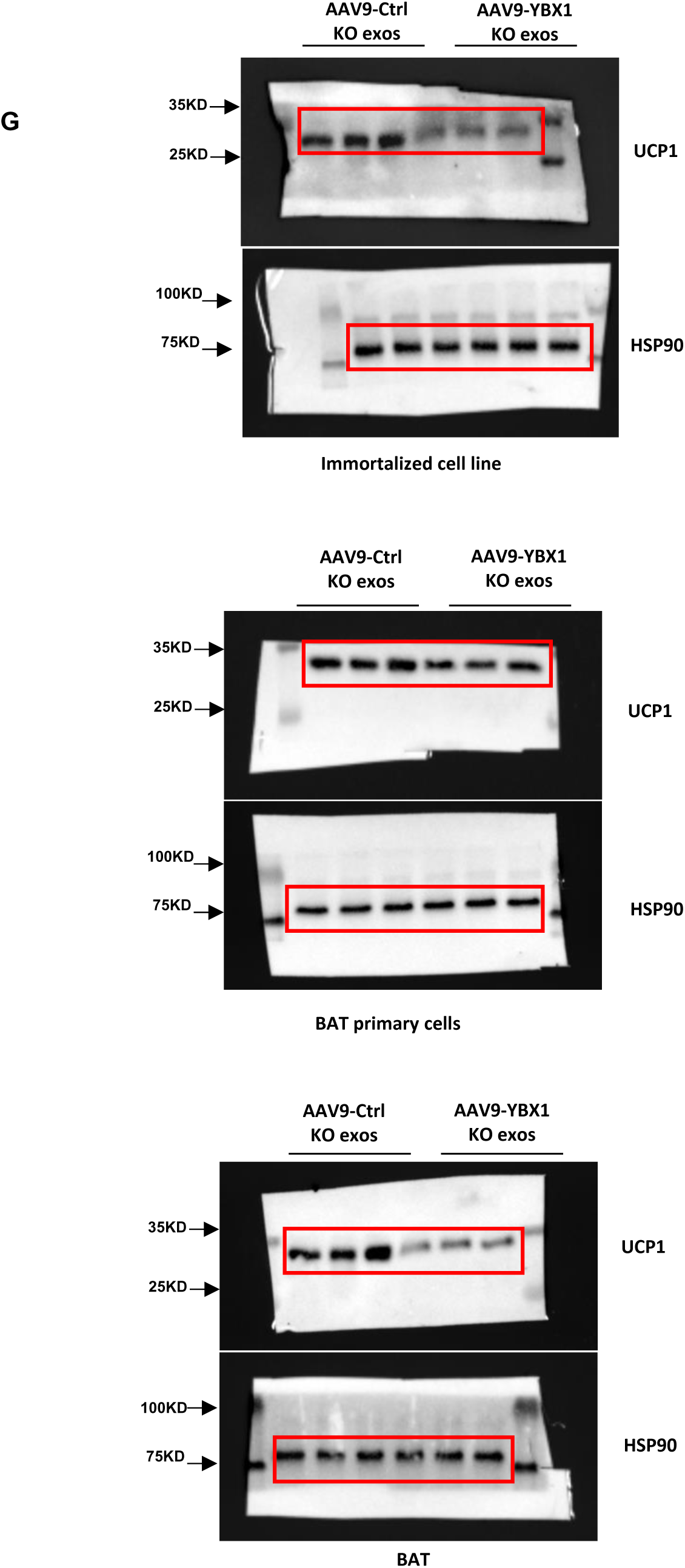
Hepatocyte exosomal miRNAs drive interorgan metabolic crosstalk. A. The in vivo protocol that used intravenously injection AAV9-Ctrl KD hepExos and AAV9-YBX1 KD hepExos. B. Body weight of mice in (A). n=5 mice each group. C. Rectal temperature was measured when mice in (A) were challenged to 4°C. n=5 mice each group. D. Fat mass weight in mice of (A). n=5 mice each group. E. Gene expression analyzed by quantitative PCR in BAT primary cells as in (A). n=3 independent experiments. F. Gene expression analyzed by quantitative PCR in BAT as in (A). n=5 independent experiments. G. UCP1 protein levels were analyzed in immortalized brown adipocytes, BAT primary cells and BAT. n=3 independent experiments. H. C57BL/6J male mice were subjected to the indirect calorimetry analysis at basal state. n=5 mice each group. I. H&E staining and protein expression of UCP1 detected by immunohistochemistry in (A) of mice from different treatment groups. Scale bar: 0.2mm for BAT.

### miR-293-5p orchestrates adipocyte lipid homeostasis

To delineate bioactive mediators underlying hepatic exosome-mediated lipid regulation, we focused on miRNA cargoes given their emerging roles in orchestrating adipose metabolism and inflammatory signaling in BAT. Small RNA sequencing of hepExos revealed miR-293-5p as the most significantly upregulated miRNA in liver , circulation, and BAT-derived exosomes following cold exposure (Fig. 3A, Table S1). Strikingly, this miRNA represented the sole candidate showing concurrent elevation across all three compartments. Quantitative RT-PCR profiling demonstrated cold-induced miR-293-5p upregulation specifically in hepatic tissue exosomes, with no significant change in BAT (Fig. 3B). This compartmentalized expression pattern strongly supports the existence of a hepatic exosomal miRNA-adipose axis mediating interorgan metabolic communication. The spatiotemporal dynamics and tissue specificity of miR-293-5p elevation prompted focused investigation into its metabolic regulatory functions.

**Figure 3.**
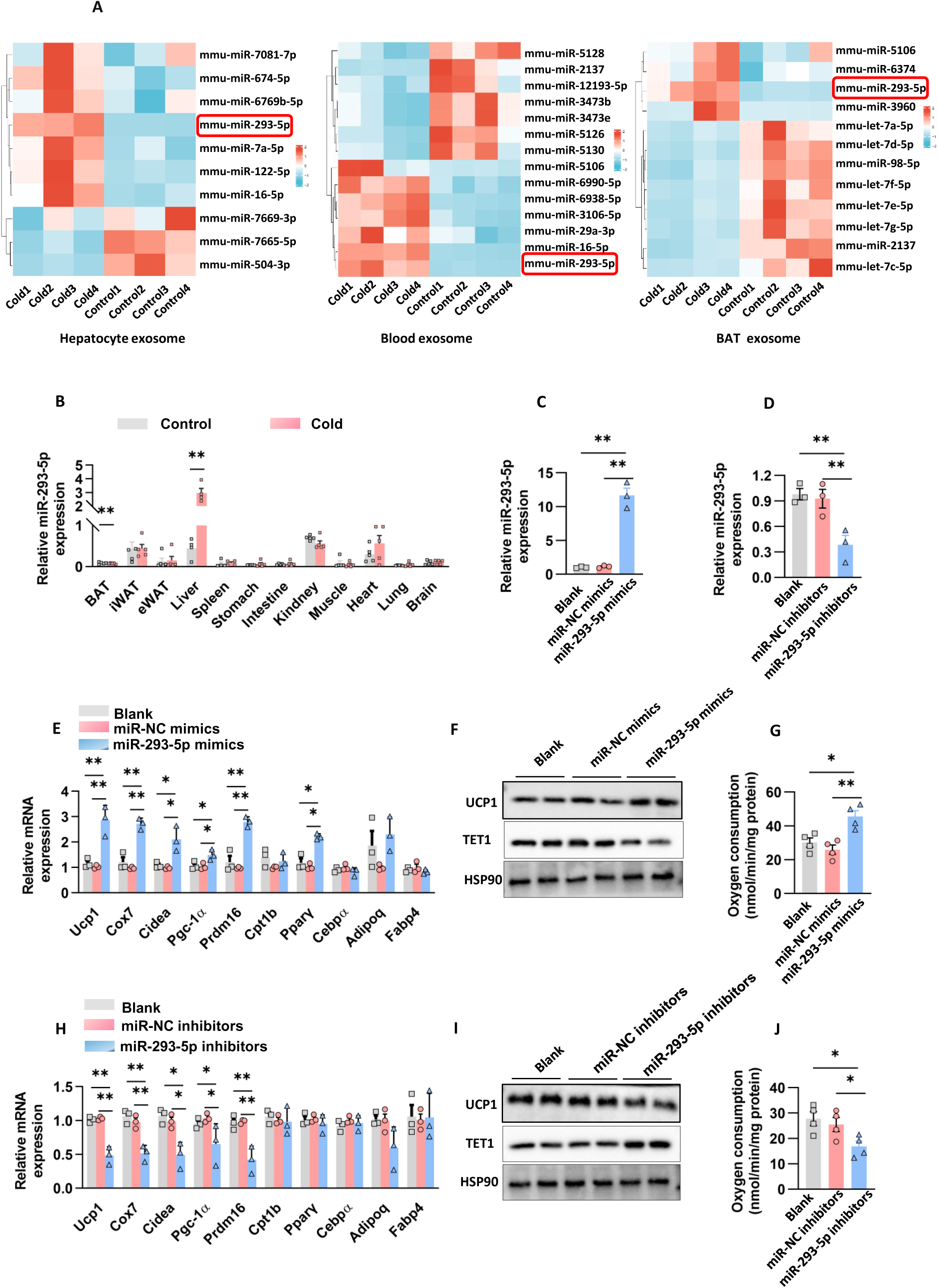
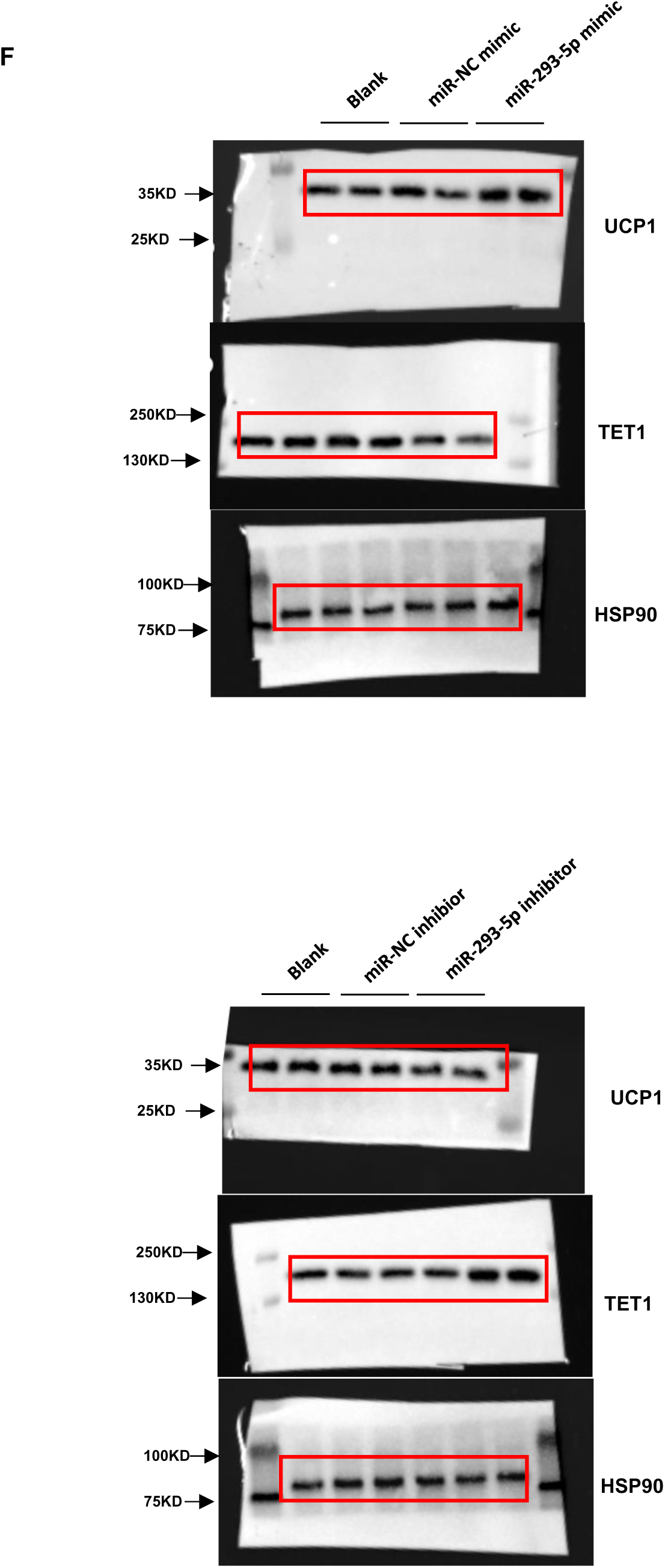
miR-293-5p orchestrates adipocyte lipid homeostasis. A. Differential expression of miRNAs from cold-exposed mice within hepatocyte, blood and BAT exosomes. B. Fold change of miR-293-5p expression level in multiple tissues after cold exposure. n=5 mice each group. C. Immortalized brown adipocytes were induced to mature adipocytes with supplementation of blank, miR-NC mimics and miR-293-5p mimics on day 3 during differentiation. Fold change of miR-293-5p expression level was analyzed on differentiation day 6. n=3 independent experiments. D. Immortalized brown adipocytes were induced to mature adipocytes with supplementation of blank, miR-NC inhibitors and miR-293-5p inhibitors on day 3 during differentiation. Fold change of miR-293-5p expression level was analyzed on differentiation day 6. n=3 independent experiments. E. Gene expression was analyzed on differentiation day 6 as in (C). n=3 independent experiments. F. UCP1 and TET1 protein levels were analyzed on day 6 as in (C). n=2 independent experiments. G. Oxygen consumption rate measured by Clark electrodes as in (C). n=3 independent experiments. H. Gene expression was analyzed on differentiation day 6 as in (D). n=3 independent experiments. I. UCP1 and TET1 protein levels were analyzed on day 6 as in (D). n=2 independent experiments. J. Oxygen consumption rate measured by Clark electrodes as in (D). n=3 independent experiments.

To functionally characterize miR-293-5p’s metabolic regulatory role, we performed gain- and loss-of-function studies during in vitro adipocyte differentiation. Transfection with miR-293-5p mimics achieved robust overexpression, while inhibitors effectively reduced endogenous levels, as quantified by RT-qPCR (Figs. 3C-D). Mimic transfection specifically upregulated thermogenic gene expression in immortalized brown adipocytes, with *Ucp1, Cox7a, Cidea, Pgc-1α,* and *Prdm16* showing significant induction, whereas lipogenic markers (*Cebpα, Adipoq, Fabp4*) and β-oxidation gene *Cpt1b* remained unaffected (Fig. 3E). Despite moderate *Pparγ* upregulation, early-stage clonal expansion was unaltered (Supp. Fig. 5A). The thermogenic enhancement was corroborated by increased UCP1 protein ( Fig. 3F) and elevation in mitochondrial oxygen consumption rate (Fig. 3G). Conversely, inhibitor treatment preserved adipogenic differentiation markers (Fig. 3H) and proliferative capacity (Supp. Fig. 5B), but severely compromised thermogenic competence- reducing UCP1 protein (Fig.3I) and suppressing thermogenic transcripts (*Ucp1,Cox7a, Cidea,Pgc-1α,Prdm16)* (Fig. 3H), ultimately decreasing OCR (Fig.3J). These complementary findings establish miR-293-5p as a critical and selective regulator of thermogenic programming in adipocytes, independent of general adipogenic differentiation processes.

### Primary hepatocyte derived exosomal miR-293-5p activates adipocyte thermogenic program

The secretion of exosomes is a tightly regulated process involving multiple steps controlled by specific molecular machinery[21]. To elucidate miR-293-5p’s regulatory role in hepExos biology, we first established a controlled secretion model using primary hepatocytes. Transfection with miR-293-5p mimics increased exosomal miR- 293-5p levels by approximately 10-fold compared to scramble controls (Fig. 4A), whereas inhibitors reduced exosomal content by 30% (Fig. 4B). Lentiviral-mediated knockout (KO-1/KO-2) and overexpression (OE-1/OE-2) systems demonstrated dose-dependent modulation of exosomal miR-293-5p levels (Figs. 4C-D), validated through nanoparticle tracking analysis and qRT-PCR.

**Figure 4.**
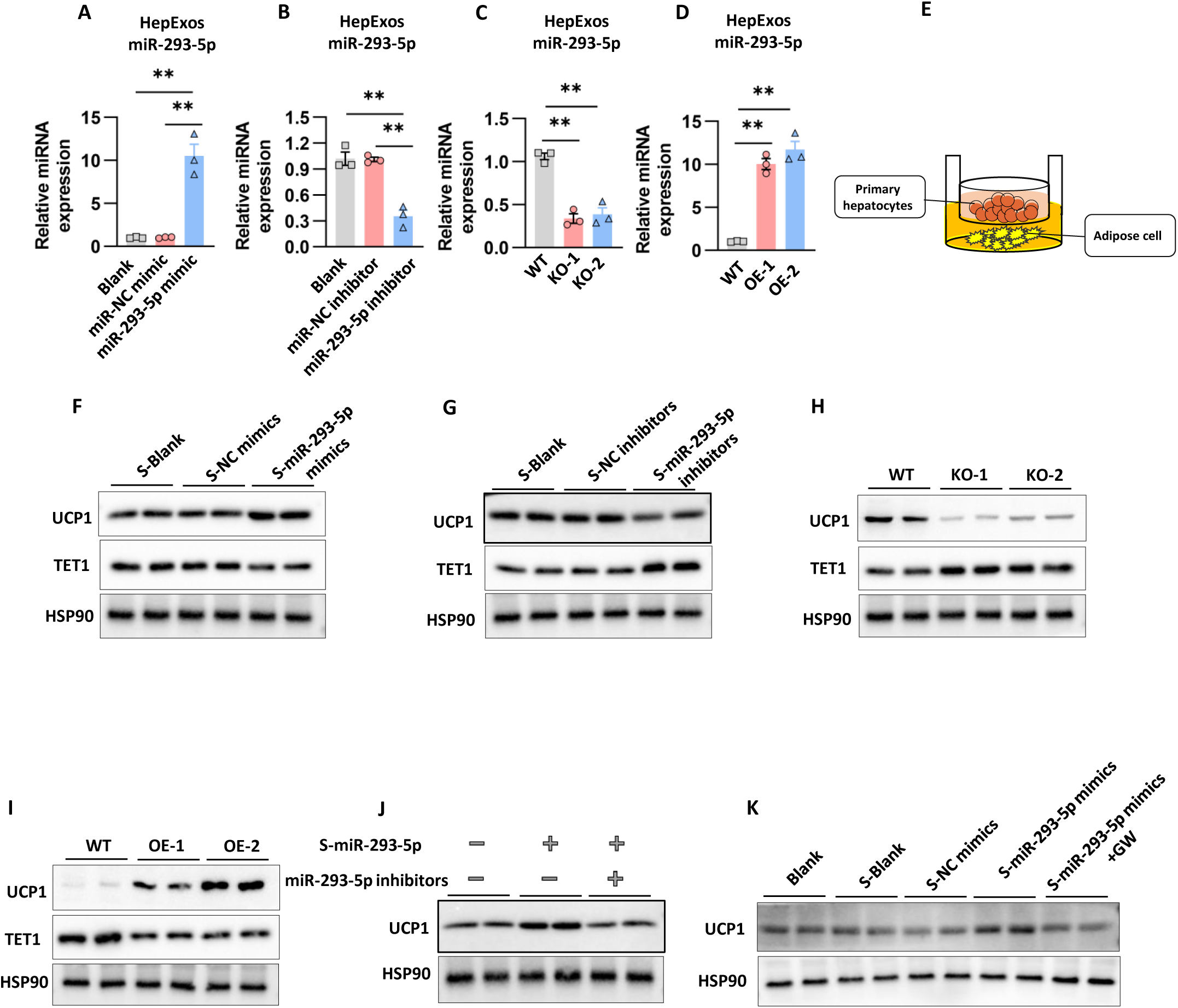
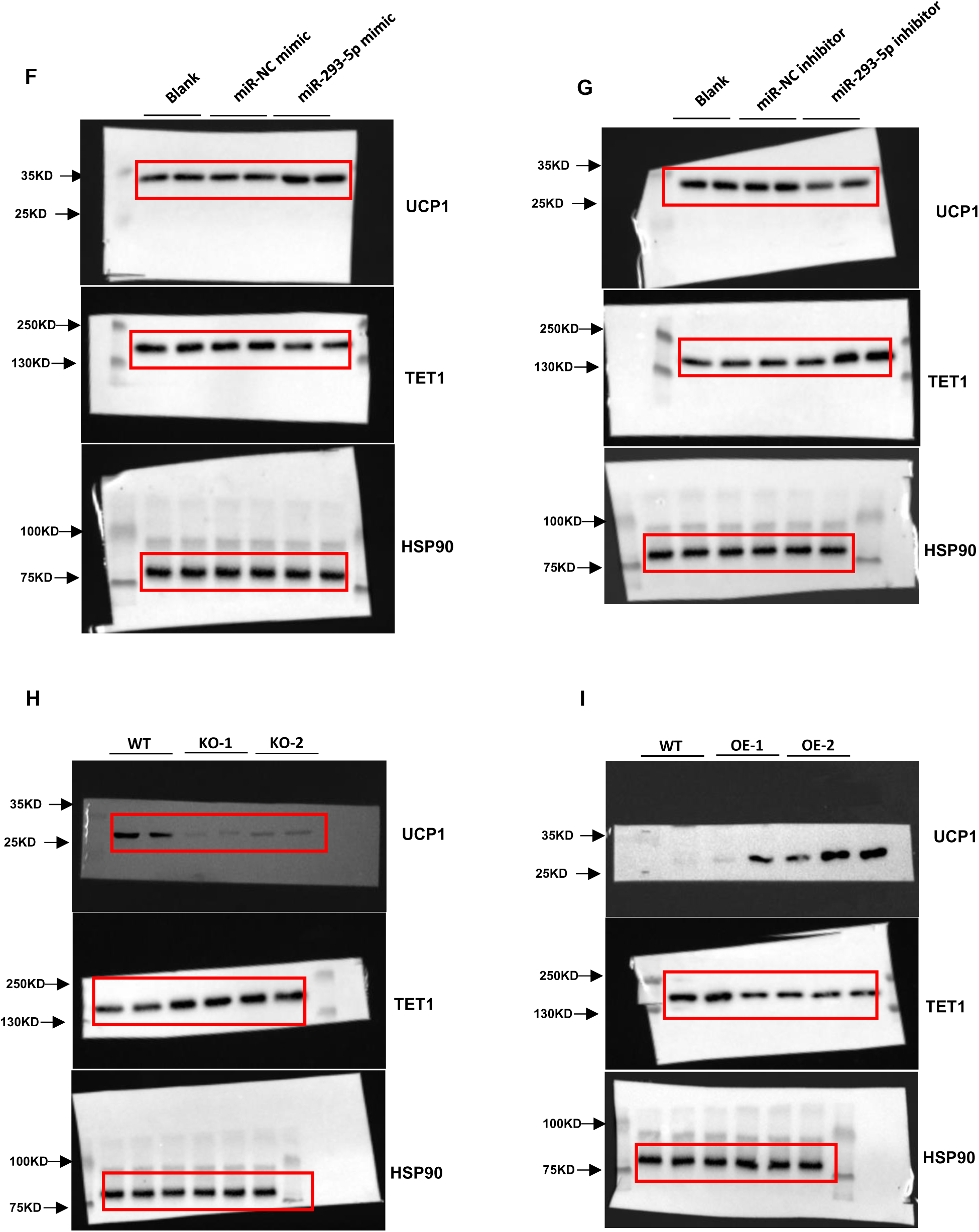
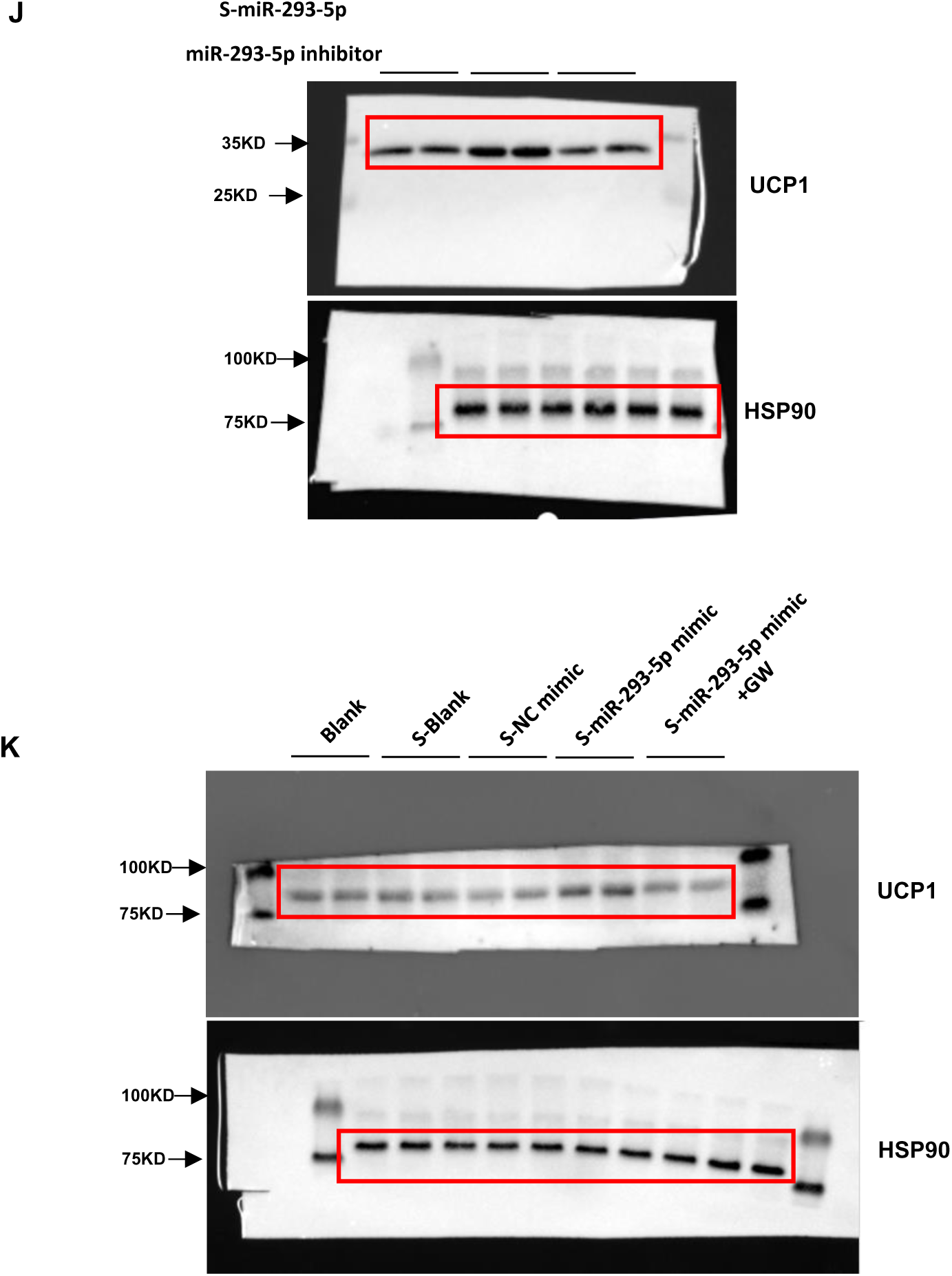
Primary hepatocyte derived exosomal miR-293-5p activates adipocyte thermogenic program. A. miR-293-5p levels in exosomes from culture medium of primary hepatocytes transfected with miR-NC mimic or miR-293-5p mimic. n=3 independent experiments. B. miR-293-5p levels in exosomes from culture medium of primary hepatocytes transfected with miR-NC inhibitor or miR-293-5p inhibitor. n=3 independent experiments. C. miR-293-5p levels in exosomes from culture medium of primary hepatocytes infected by lentiviral shRNAs against miR-293-5p. n=3 independent experiments. D. miR-293-5p levels in exosomes from culture medium of primary hepatocytes infected by lentiviral overexpressed miR-293-5p. n=3 independent experiments. E.Schematic of the transwell coculture chamber. F,G. Primary hepatocytes were transfected with miR-293-5p mimics or inhibitors and the control. The indicated culture medium was then harvested to further studies. Western blot analysis of UCP1, and TET1 expression in immortalized brown adipocytes incubated with conditioned medium. n=2 independent experiments. H,I. Primary hepatocytes were infected by lentiviral shRNAs against miR-293- 5p or infected by lentiviral overexpressed miR-293-5p. The indicated culture medium was then harvested to further studies. Western blot analysis of UCP1, and TET1 expression in immortalized brown adipocytes incubated with conditioned medium. n=2 independent experiments. J. Western blot analysis of UCP1 expression in immortalized brown adipocytes transfected with miR-293-5p inhibitor for 24 h, and then incubated with conditioned medium from primary hepatocytes overexpressing miR-293-5p for 48 h. n=2 independent experiments. K. Western blot analysis of UCP1 expression in immortalized brown adipocytes incubated with conditioned medium from the indicated primary hepatocytes. n=2 independent experiments.

In a transwell coculture system modeling hepatic-adipose crosstalk (Fig. 4E), miR-293-5p-enriched hepExos from mimic-transfected hepatocytes induced UCP1 upregulation in brown adipocytes (Fig. 4F). Conversely, inhibitor-treated hepatocytes reduced adipocyte UCP1 (Fig. 4G). Genetic perturbation via lentiviral knockout (Fig. 4C) recapitulated this suppression, decreasing UCP1 expression of controls (Fig. 4H), while overexpression systems (OE-1/OE-2) enhanced UCP1 levels (Fig. 4I). These coordinated findings across pharmacological and genetic approaches confirm hepatocyte-derived exosomal miR-293-5p as a potent regulator of adipocyte thermogenesis, operating through exosome-dependent intercellular signaling.

To further validate the hepExos-miR-293-5p in thermogenic regulation, we conducted a series of functional rescue experiments. Prior to exposure to conditioned media from miR-293-5p-overexpressing primary hepatocytes, adipocytes were pre-incubated with a specific miR-293-5p inhibitor. Remarkably, this pharmacological intervention completely abolished the upregulation of UCP1 expression induced by hepatocyte-derived factors (Fig. 4J), demonstrating the miRNA’s essential role in mediating this regulatory effect. To specifically address the involvement of extracellular vesicles (EVs), we employed GW4869 - a well-characterized inhibitor of exosome biogenesis and secretion. Pharmacological blockade of EV production effectively attenuated the thermogenic activation in adipocytes treated with media from miR-293-5p-overexpressing hepatocytes (Fig.4K). These complementary experimental approaches provide conclusive evidence that hepExos serve as critical vehicles for miR-293-5p delivery, and that this intercellular communication mechanism is indispensable for initiating thermogenic programming in adipocytes.

### Injecting agomir miR-293-5p activates thermogenesis in BAT

To investigate the in vivo necessity of miR-293-5p in thermogenic regulation, wild-type mice received weekly intravenous injections of agomir miR-293-5p or control miR-NC once a week for 6 weeks (Fig. 5A). Pharmacological activation achieved 3-fold elevation of miR-293-5p in BAT versus controls (Fig. 5B), without altering basal metabolic parameters including food/water consumption, body weight progression, or serum lipid profiles (Supp. Figs. 6A-C). Notably, miR-293-5p potentiation induced progressive metabolic adaptations characterized by: enhanced cold tolerance, delayed-onset body weight, reduction primarily through eWAT mass depletion, and increased BAT OCR (Figs. 5C-F).

**Figure 5.**
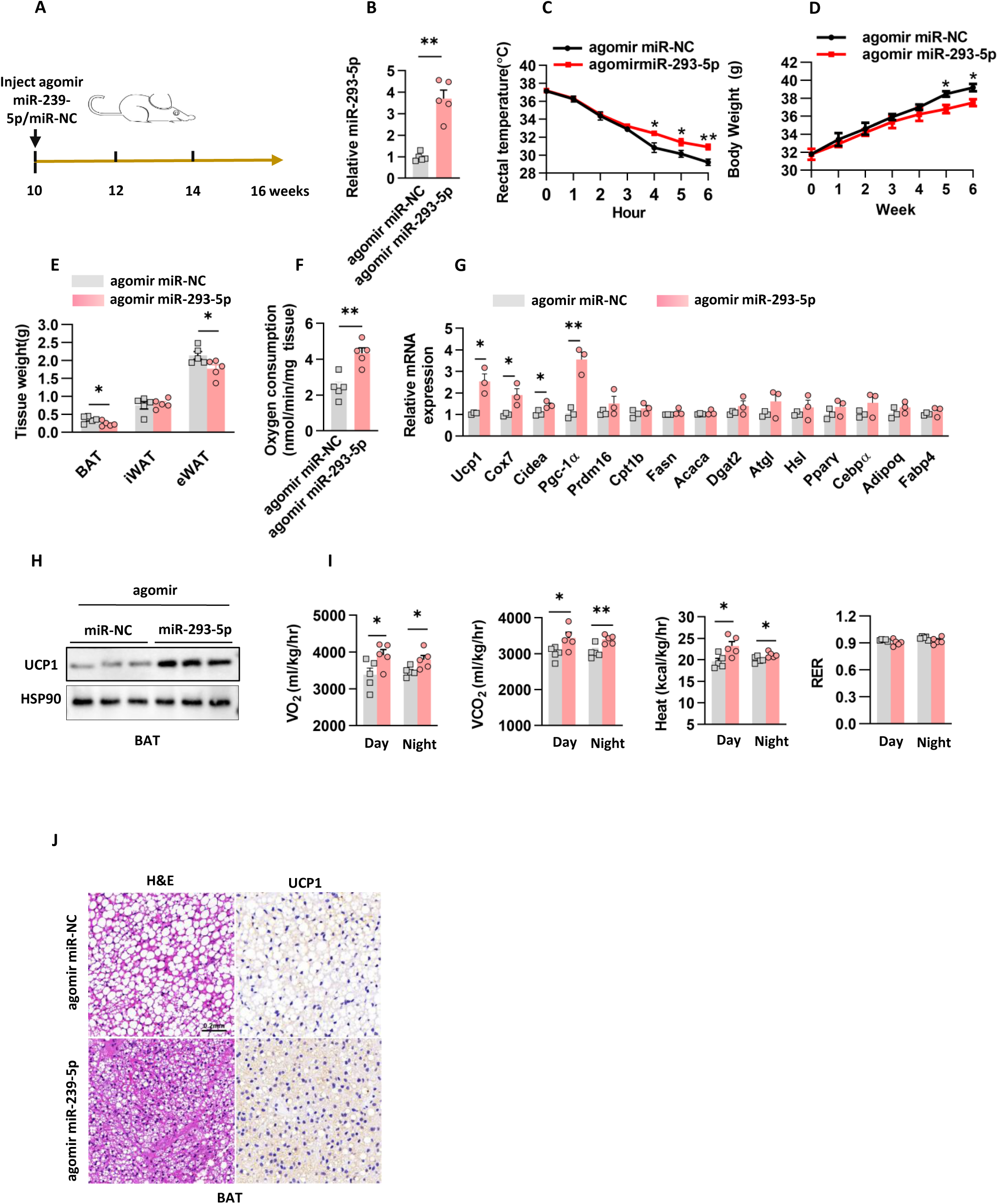
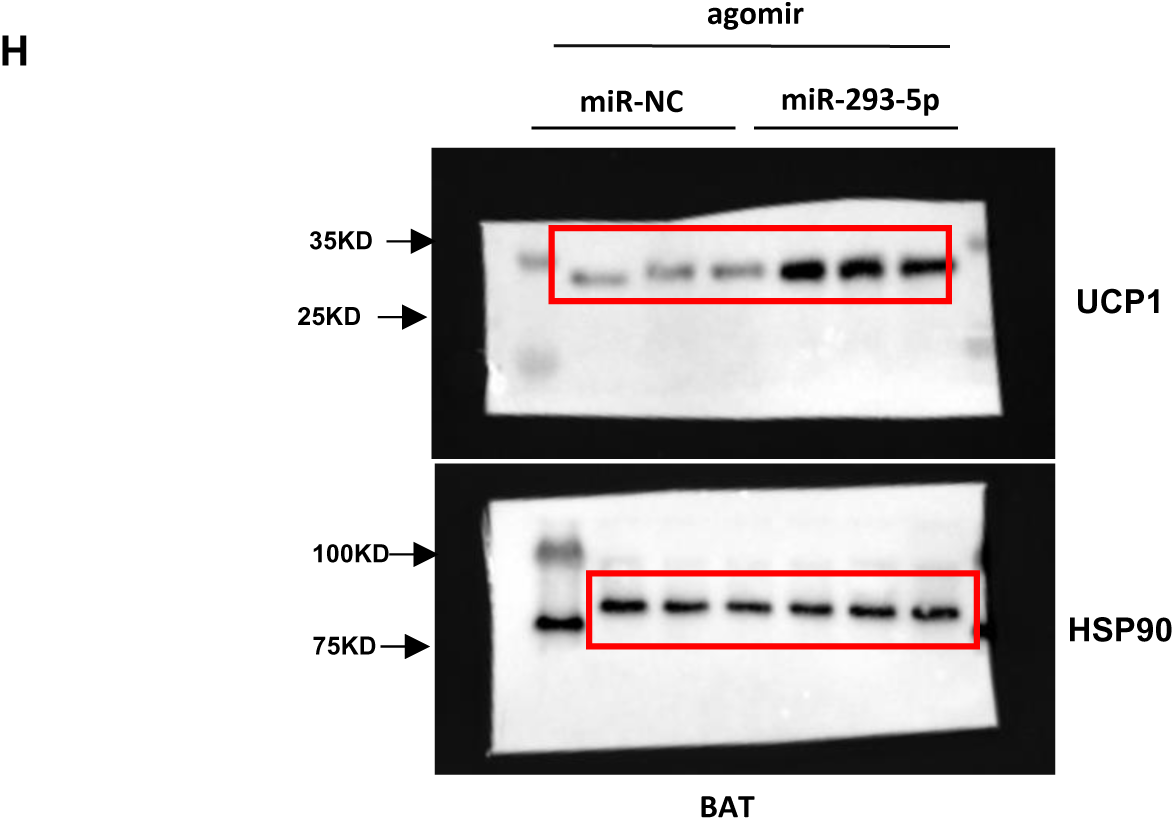
Injecting agomir miR-293-5p activates thermogenesis in BAT. A. The in vivo protocol that used tail vein injection agomir miR-293-5p or miR-NC. B. miR-293-5p levels in BAT of mice in (A). n=5 mice each group. C. Rectal temperature was measured when mice in (A) were challenged to 4°C. n=5 mice each group. D. Body weight of mice in (A). n=5 mice each group. E. Fat mass weight in mice of (A). n=5 mice each group. F. Oxygen consumption rate measured by Clark electrodes for BAT as in (A). n=5 independent experiments. G. Gene expression analyzed by quantitative PCR in BAT as in (A). n=3 independent experiments. H. UCP1 protein levels were analyzed in BAT. n=3 independent experiments. I. C57BL/6J male mice were subjected to the indirect calorimetry analysis at basal state. n=5 mice each group. J. H&E staining and protein expression of UCP1 detected by immunohistochemistry in (A) of mice from different treatment groups. Scale bar: 0.2mm for BAT.

Molecular analyses revealed coordinated upregulation of thermogenic effectors in BAT, with UCP1 protein levels increasing accompanied by transcriptional activation of mitochondrial uncoupling genes (*Ucp1, Cox7a, Cidea, Pgc-1α*)(Figs. 5G,H). Whole-body calorimetry demonstrated elevated energy expenditure paralleled by BAT morphological remodeling - reduction in lipid droplet size and dense UCP1⁺ adipocyte accumulation (Figs. 5I-J). The tissue-specific nature of this response was evidenced by minimal gene expression changes in iWAT. These multimodal data establish miR-293-5p as a critical enhancer of BAT-driven thermogenesis through mitochondrial activation and adipose tissue remodeling.

### miR-293-5p-targeted therapy attenuated obesity progression in mice

Previous studies have established that obesity-induced dysfunction in β3- adrenergic signaling pathways compromises the thermogenic capacity of brown and beige adipocytes [23,24]. To explore the therapeutic potential of miR-293-5p in restoring adipose tissue responsiveness under obese conditions, we conducted a longitudinal intervention study in diet-induced obese mice. During 10 weeks of high-fat diet feeding, animals received systemic administration of either agomir miR-293- 5p or miR-NC through intravenous delivery (Fig. 6A). While no intergroup differences were observed in caloric intake or hydration status (Supp. Figs. 7A,B), a progressive divergence in body weight trajectories emerged, with agomir miR-293- 5p-treated mice demonstrating significant weight reduction from week 7 onward (Fig. 6B). Body composition analysis revealed this anti-obesity effect principally originated from marked decreases in both inguinal and eWAT depots (Fig. 6C). The metabolic improvements extended beyond adiposity measures. miR-293-5p-enhanced mice maintained superior thermoregulatory capacity during cold challenge, as evidenced by sustained higher core body temperatures (Fig. 6D). Circulating metabolic parameters showed comprehensive amelioration, with significant reductions in triglyceride levels total cholesterol, and free fatty acids compared to controls (Figs. 6E-G). Functional assessments demonstrated enhanced whole-body metabolic flexibility, including improved glucose disposal rates and heightened insulin sensitivity (Figs. 6H,I). Mechanistic investigations revealed elevated UCP1 protein expression in BAT, though this trend did not reach statistical significance in iWAT (Fig 6J; Supp. Figs. 7D). Indirect calorimetry measurements corroborated these findings, showing increased heat production accompanied by elevated VO2 and VCO2 in the treatment group (Fig 6K). Histopathological evaluation of BAT demonstrated reduced lipid droplet size and enhanced multilocular morphology, indicative of activated thermogenic potential (Fig 6L). Collectively, these multimodal data establish miR- 293-5p administration as an effective strategy for counteracting diet-induced metabolic dysregulation through coordinated enhancement of adipose tissue functionality and systemic energy expenditure.

**Figure 6.**
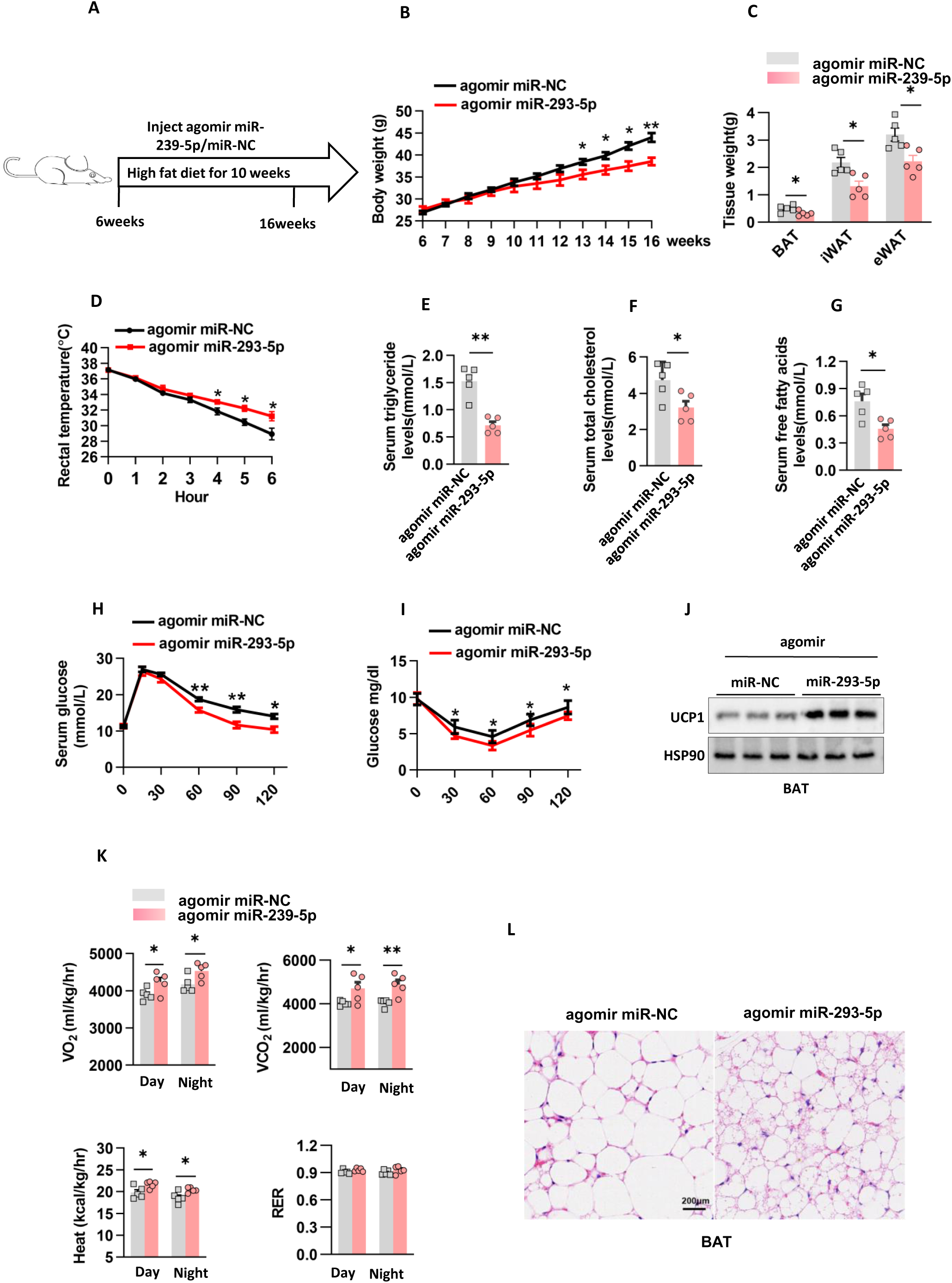
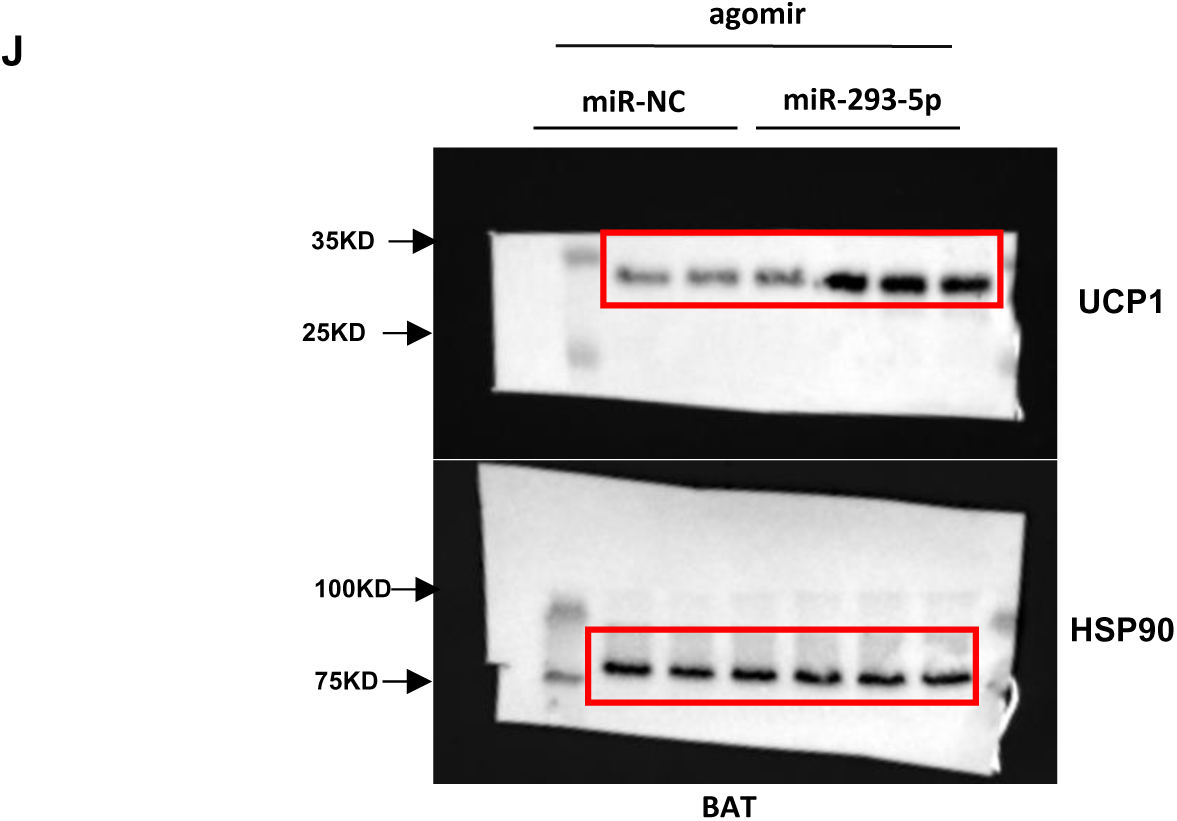
miR-293-5p-targeted therapy attenuated obesity progression in mice. A. C57BL/6J male mice fed on a high fat diet protocol that used tail vein injection agomir miR-293-5p or miR-NC. B. Body weight of mice in (A). n=5 mice each group. C. Fat mass weight in mice of (A). n=5 mice each group. D. Rectal temperature was measured when mice in (A) were challenged to 4°C. n=5 mice each group. E-G. Serum triglyceride (E), serum total cholesterol (F) and serum free fatty acids (G) concentration was assayed in mice of (A). n=5 mice each group. H. Glucose tolerance test was performed after high-fat diet feeding in mice of (A). n=5 mice each group. I. Insulin tolerant test was performed after high-fat diet feeding in mice of (A). n=5 mice each group. J. UCP1 protein levels were analyzed in BAT. n=3 independent experiments. K. C57BL/6J male mice were subjected to the indirect calorimetry analysis at basal state. n=5 mice each group. L. H&E staining in (A) of mice from different treatment groups. Scale bar: 0.2mm for BAT.

### Tet1 (Ten-eleven-translocation 1) serves as a direct miR-293-5p target

MicroRNAs execute post-transcriptional regulation through complementary base pairing between their seed sequences and target mRNAs, leading to translational inhibition or transcript degradation. To systematically identify miR-293-5p targets, we integrated computational predictions from three algorithms (TargetScan, miRmap, miRDB) with transcriptome profiling (Fig. 7A). This multi-platform approach identified Tet1 (Ten-eleven translocation methylcytosine dioxygenase 1) as a high-confidence candidate, showing reduction in mRNA levels (Figs. 7B) and decrease in protein expression following miR-293-5p mimic transfection in adipocytes (Fig. 3F). The regulatory specificity was confirmed through antisense inhibition experiments, where miR-293-5p inhibitors elevated TET1 protein levels (Fig. 3I). Cross-tissue validation in hepatocyte-adipocyte coculture systems revealed cell-nonautonomous regulation: adipocytes exposed to miR-293-5p-overexpressing hepatocytes exhibited lower TET1 protein level (Figs. 4F,I), whereas miR-293-5p knockout or inhibitor-treated systems increased TET1 expression (Figs. 4G,H). Mechanistically, luciferase reporter assays in 293T cells demonstrated direct targeting, with wild-type TET1 3’UTR activity suppressed by 45% through miR-293-5p binding - an effect abolished by seed sequence mutations (Fig. 7C,D). To establish direct miR-293-5p/TET1 targeting, we transfected miR-293-5p mimics and TET1 mRNA in 293T cells. The transfection of miR-293-5p mimics significantly reduced TET1 protein levels (Fig. 7E). However, this effect was not observed with mutant TET1 protein (Fig. 7F). Additionally, using miR-293-5p inhibitors to counteract the effect of the miR-293-5p mimics restored TET1 protein expression (Fig. 7G). This dose-responsive, sequence-specific regulatory paradigm - combining gain-of-function suppression, loss-of-function rescue, and binding site mutagenesis - definitively characterizes TET1 as a bonafide miR-293-5p target governed by evolutionarily conserved seed-pairing interactions.

**Figure 7.**
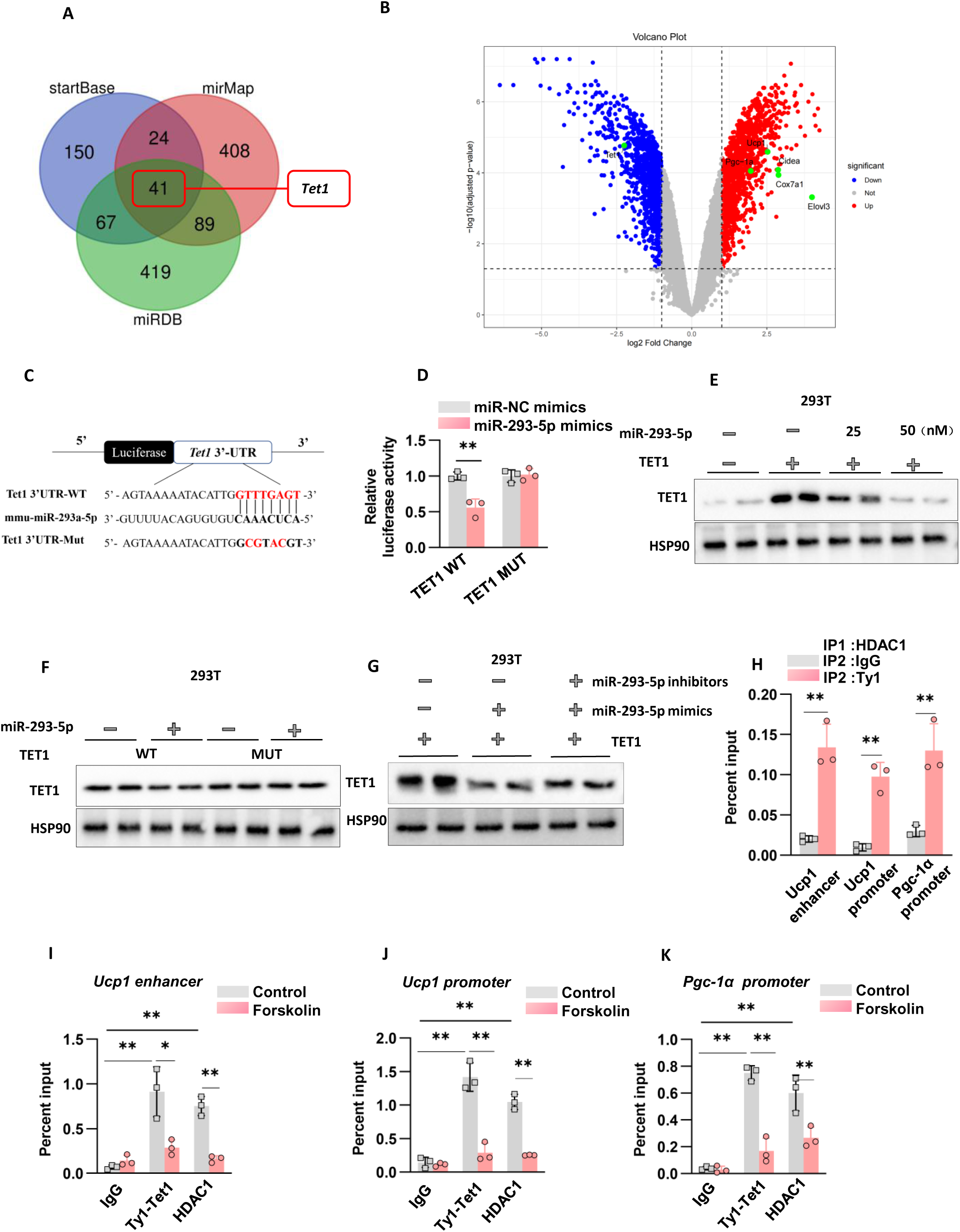
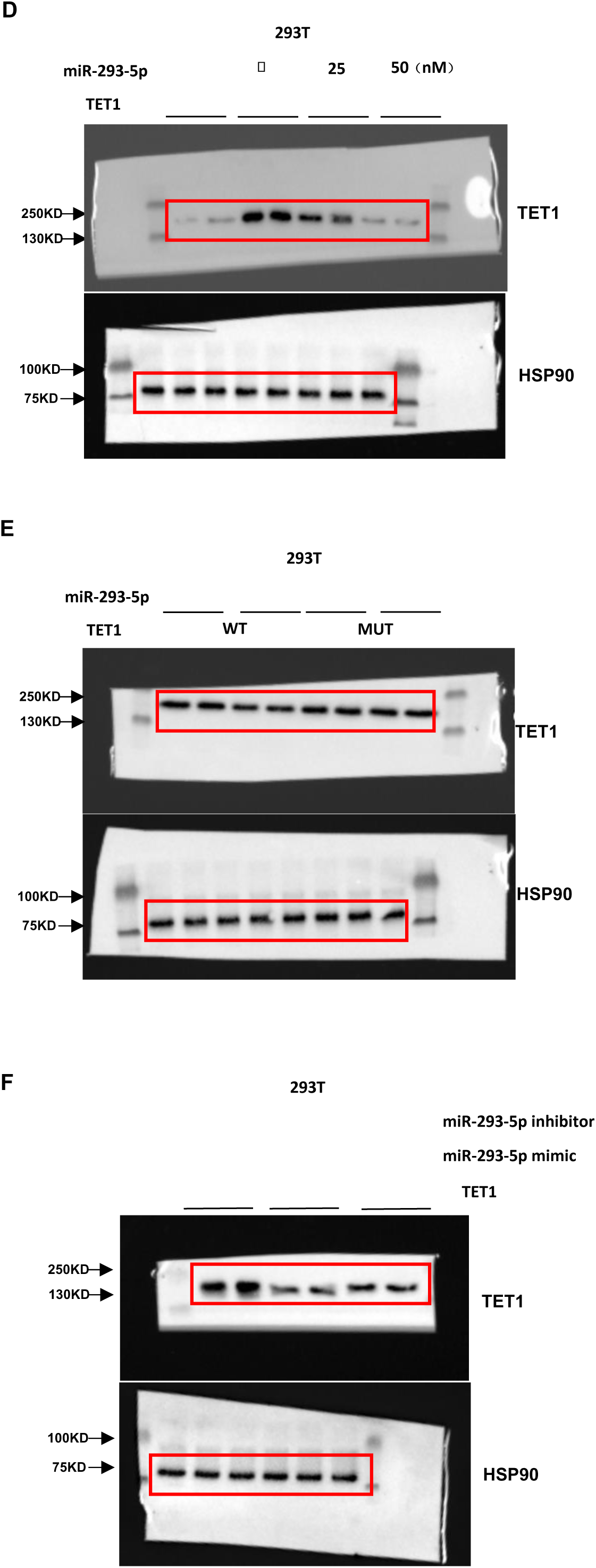
Tet1 (Ten-eleven-translocation 1) serves as a direct miR-293-5p target. A. Venn diagram depicted the target gene prediction algorithm (startBase, mirMap and miRDB) to identify theoretical target genes of miR-293-5p. B. The miRNA-seq results of primary brown fat cells treated by control hepExos and 3d cold hepExos. n=4 independent experiments. C. R-293-5p binding sites on the Tet1 3’UTRs. D. Results of the luciferase reporter assay of miR-293-5p and Tet1 in 293T cells cotransfected with WT or mutant mouse Tet1. n=3 independent experiments. E. Western blot analysis of Tet1 and Myc protein levels in response to the miR-293-5p mimics in 293T cells cotransfected with WT Tet1. F. Western blot analysis of Tet1 and Myc protein levels in response to the miR-293-5p mimics in 293T cells cotransfected with WT or mutant Tet1. G. Western blot analysis of Tet1 protein level in response to the miR-293-5p mimics and miR-293-5p mimics plus inhibitors in 293T cells cotransfected with WT Tet1. H.ChIP-reChIP qPCR analysis was performed on beige adipocytes that were transduced with Ty1-Tet1 WT or GFP using HDAC1 (1st IP), Ty1 (2nd IP), or IgG as a control. The enrichment efficiency is presented as a percent input at the indicated binding sites.n=3 independent experiments. I–K Ty1 and HDAC1 ChIP-qPCR analysis was performed in cells that express Ty1- Tet1 WT with and without forskolin stimulation.n=3 independent experiments.

Emerging evidence suggests TET1 exerts transcriptional repression through cooperative interactions with epigenetic regulators, including Polycomb Repressive Complex 2 (PRC2), SIN3A, and histone deacetylases (HDACs) across various cellular contexts. Building on our prior discovery that HDAC1 inhibition (genetic or pharmacological) robustly enhances UCP1/PGC-1α expression and oxidative metabolism[25], we hypothesized functional interplay between TET1 and HDAC1 in thermogenic gene silencing. To test this, we established a Ty1 epitope-tagged TET1 overexpression system, circumventing limitations of endogenous TET1 antibody specificity. Sequential chromatin immunoprecipitation (ChIP-reChIP) revealed significant co-occupancy of TET1 and HDAC1 at target loci (Fig. 7H). Crucially, this co-localization exhibited dynamic responsiveness to physiological stimuli—forskolin treatment markedly reduced both TET1 and HDAC1 occupancy at these genomic regions (Figs.7I–K), mirroring the known disassembly of repressive complexes during β-adrenergic receptor-activated thermogenesis.

### Overexpression of *Tet1* inhibits BAT thermogenic program in mice

To dissect TET1’s functional hierarchy within miR-293-5p-regulated thermogenic pathways, we engineered a BAT-specific TET1 gain-of-function model through adenoviral vector delivery (pAd-GFP vs. pAd-TET1) combined with agomir miR-293-5p (Figure 8A). TET1 overexpression exerted potent suppression of brown adipocyte activation, reducing UCP1 protein expression and downregulating core thermogenic regulators including *Ucp1, Cox7a , Cideα, Pgc-1α* and *Prdm16* versus GFP controls (Figs. 8B,C). Functional consequences manifested as impaired cold tolerance and reduction in mitochondrial oxygen consumption rate during cold exposure - phenotypes partly recoverable to agomir miR-293-5p (Figs. 8D,E). Histomorphometric analysis revealed TET1’s dual lipid-metabolic impact: adipocytes from miR-293-5p-treated mice displayed smaller lipid droplets alongside elevated UCP1 immunoreactivity, contrasting with TET1-overexpressing counterparts showing hypertrophic lipid accumulation and depleted UCP1 signals (Fig. 8F). This epistatic relationship demonstrates TET1 operates downstream of miR-293-5p in a non-compensatory regulatory node, where forced TET1 expression dominantly suppresses BAT activation through epigenetic silencing of thermogenic enhancers.

**Figure 8.**
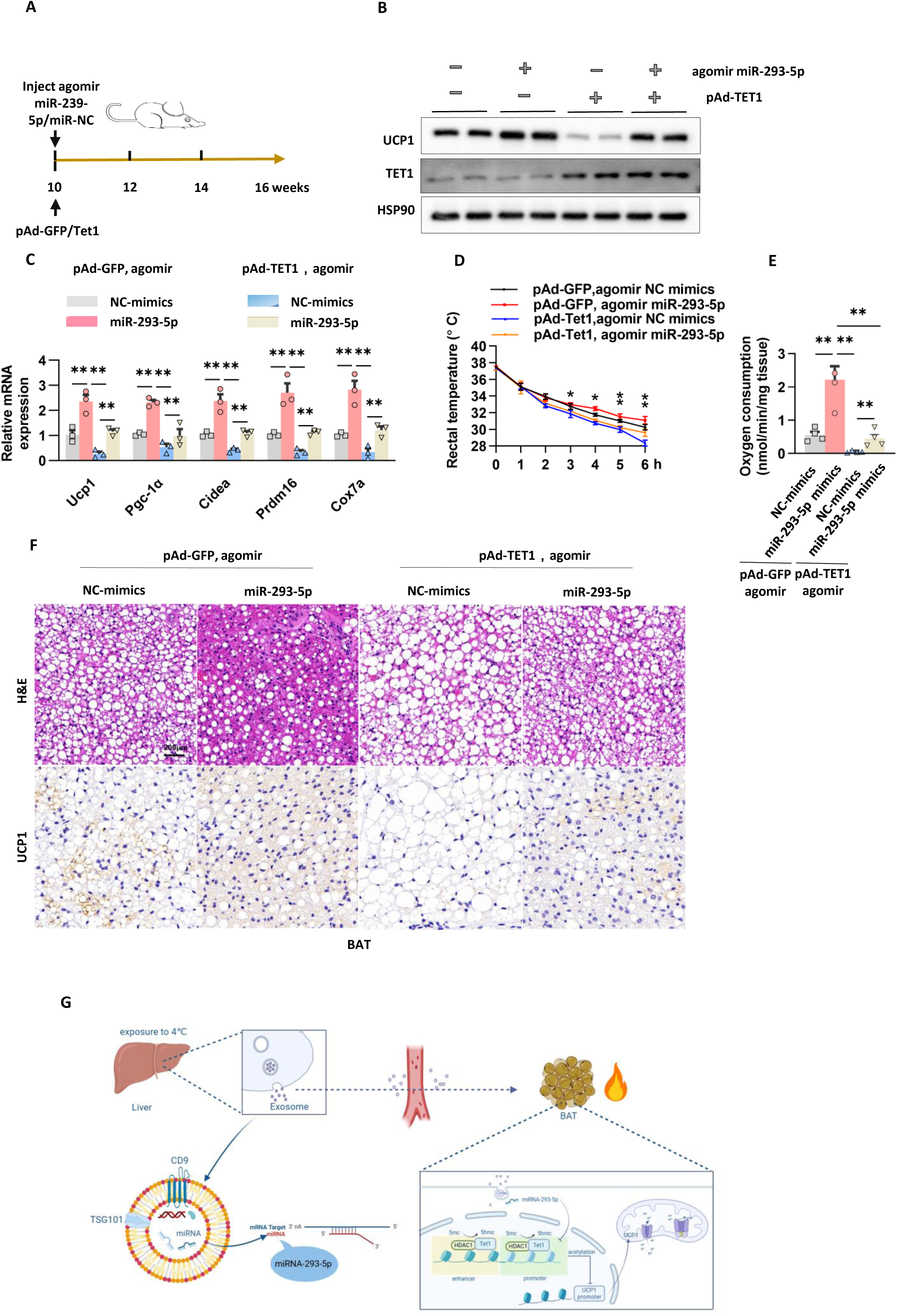
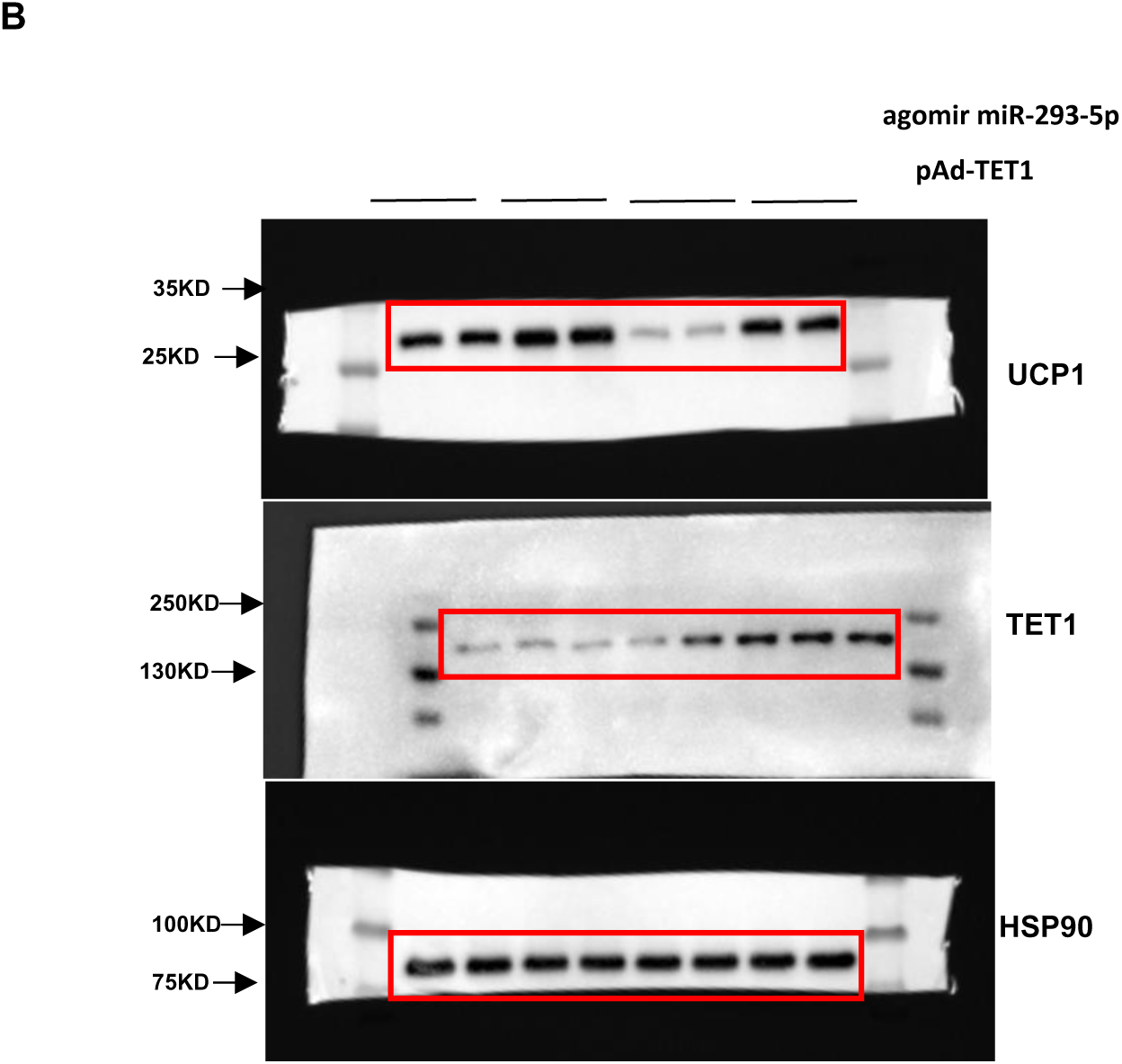
Overexpression of *Tet1* inhibits BAT thermogenic program in mice. A. Protocol of the pAd-Tet1 and agomir miR-293-5p/NC injection in mice. B. Western blot analysis of Tet1 and UCP1 protein level in response to the agomir miR-293-5p/NC injection in mice cotransfected with WT mouse pAd-TET1. C. Gene expression analyzed by quantitative PCR for BAT as in (A). n=3 independent experiments. D. Core body temperature of mice from different treatment groups. n=5 independent experiments. E. Oxygen consumption rate measured by Clark electrodes for BAT as in (A). n=4 independent experiments. F. H&E staining and protein expression of UCP1 detected by immunohistochemistry in the BAT of mice from different treatment groups. Scale bar: 0.2mm. G. Cold exposure activates the YBX1-dependent exosomal miRNA sorting machinery in hepatocytes, driving selective enrichment of miR-293-5p within hepatic exosomes. These miRNA-loaded vesicles are subsequently transported via systemic circulation to adipose depots, where miR-293-5p directly targets Tet1 mRNA through complementary 3’UTR binding, thereby alleviating Tet1-mediated epigenetic repression of thermogenic genes. This interorgan signaling cascade culminates in enhanced mitochondrial uncoupling via UCP1 upregulation and amplified oxidative phosphorylation capacity, ultimately activating BAT-mediated energy expenditure.

## Discussion

Our investigation elucidates hepExos as critical mediators of cold-adaptive thermogenesis through systemic miRNA trafficking. Three-day cold exposure triggered increase in hepatic exosome secretion, with these thermogenically primed vesicles demonstrating direct BAT activation capacity through UCP1 upregulation and mitochondrial uncoupling enhancement. Cross-tissue miRNA profiling coupled with functional screening identified miR-293-5p as the central regulator, showing elevation in hepatocytes, circulating exosomes, and BAT under cold stress. Therapeutic miR-293-5p agonism amplified cold-induced thermogenesis, increasing BAT UCP1 protein expression while reducing visceral adiposity in diet-induced obese mice. This interorgan communication axis, operating through cold-inducible exosomal miRNA transfer, demonstrates therapeutic potential for metabolic syndrome via targeted potentiation of adaptive thermogenesis. The identification of miR-293-5p as both biomarker and functional effector in this pathway provides dual diagnostic and interventional opportunities for obesity management.

Exosomes are formed through the endocytosis-mediated invagination of the plasma membrane. Extensive evidence supports that exosomes are crucial mediators of intracellular communication and intertissue crosstalk by delivering cargo to target cells and interacting with cell surface receptors[26,27]. Exosomes contain various molecules, including proteins, mRNA, miRNA, and lncRNA, with miRNAs being critical for the majority of their biological effects [28,29]. Adipocyte-derived exosomal miRNAs are known to coordinate communication between adipocytes and other key metabolic organs [30,31]. However, the metabolic regulatory potential of hepatocyte-derived exosomal miRNAs in adipose biology remains largely uncharted. To address this knowledge gap, we engineered hepatic-specific YBX1 KD models, given YBX1’s established role in miRNA loading into exosomes. Hepatic YBX1 deficiency decreased exosomal miRNA content, concomitant with impaired cold-induced BAT activation. Small RNA sequencing of hepatocyte-derived exosomes from cold-adapted wild-type mice identified miR-293-5p as the predominant cold-responsive species. Notably, miR-293-5p emerged as the only miRNA exhibiting multi-compartment synchronization, with cold-induced upregulation in hepatocyte-derived exosomes , circulating exosomes, and BAT exosomes. qRT-PCR validation revealed strict spatial regulation - hepatic exosomal miR-293-5p increased versus controls, while BAT-resident miR-293-5p remained unchanged. This compartment-specific amplification, confirms a cold-inducible secretory mechanism rather than autonomous adipose production. The observed endocrine (hepatic secretion → systemic circulation → BAT uptake) provides mechanistic validation of an exosome- facilitated liver-adipose signaling axis governing adaptive thermogenesis.

Functional validation experiments established miR-293-5p as the principal effector of hepatic exosome-mediated thermogenesis. In vitro administration of miR- 293-5p mimics to differentiated 3T3-L1 adipocytes specifically upregulated thermogenic markers without altering adipogenic differentiation markers, confirming pathway-specific activation. The hepatic exosomal origin of this regulation was evidenced by miR-293-5p’s predominance within hepatocyte-derived EVs. Systemic delivery of agomir miR-293-5p in diet-induced obese mice reversed metabolic dysfunction, enhancing glucose disposal and insulin sensitivity. MiR-293-5p’s dual metabolic effects stem from BAT-selective thermogenic reprogramming and WAT browning, synergistically enhancing systemic energy expenditure. These coordinated multi-tissue adaptations confirm miR-293-5p’s role as a master coordinator of interorgan metabolic homeostasis through exosome-mediated epigenetic regulation.

The Ten-Eleven Translocation (TET) dioxygenase family—comprising TET1, TET2, and TET3—orchestrate epigenetic regulation through iterative oxidation of 5- methylcytosine (5mC) to 5-hydroxymethylcytosine (5hmC) and subsequent derivatives, dynamically modulating DNA methylation landscapes [32,33]. Beyond their established roles in developmental processes, TET dysregulation drives pathogenesis in hematologic malignancies[34,35]. Our work reveals a novel metabolic regulatory axis: miR-293-5p directly targets TET1, suppressing its expression by 45% in brown adipocytes, as demonstrated through dual-luciferase assays with wild-type versus seed sequence-mutated TET1 3’UTR constructs. This miRNA-mediated repression is fully reversible, with locked nucleic acid (LNA) inhibitors restoring. Mechanistically, TET1 coordinates with HDAC1 to enforce epigenetic silencing at thermogenic loci. Adipose-specific TET1 ablation in mice increases energy expenditure and confers metabolic resilience, reducing HFD-induced weight gain and improving glucose tolerance[36]. Paradoxically, while TET1 supports brown adipogenesis via Prdm16 promoter demethylation [37], it concurrently represses mature adipocyte thermogenesis through PPARγ deacetylation . This duality mirrors AMPKα1-deficient models where α-KG depletion impairs TET-mediated Prdm16 activation, causing BAT dysfunction [38]. The bimodal regulatory paradigm—pro-adipogenic yet anti-thermogenic—parallels Zfp423’s dual functionality. While essential for white adipocyte commitment[39], ZFP423 simultaneously restricts beigeing by sequestering EBF2 coactivators and reducing PPARγ occupancy at UCP1 enhancers[40]. Similarly, TET1’s context-dependent effects—facilitating adipocyte differentiation while constraining thermogenic capacity—highlight epigenetic "brakes" that could be therapeutically modulated to uncouple adipogenesis from metabolic dysfunction.

Limitations exist for this study. Firstly, liver tissue contains various cell types, including hepatocytes, immune cells, and endothelial cells. This research focuses solely on exosomes originated from hepatocytes, although the it accounts for a large proportion of extracellular vesicles entering blood from hepatocytes. Future studies should focus on the use of exosomes isolated from liver non-parenchymal cells (NPC) to rule out implications for this study. Secondly, we focused our study on the miRNAs and identified miR-293-5p as the critical microRNAs that affect BAT metabolism by comparing the miRNA profiles of 3-day cold exposured mice. Moreover, exosomal cargoes other than miRNAs may also mediate the effect of hepExos on lipid metabolism. We conducted further experiments to profile the protein contents of hepExos. We pinpointed a few protein candidates whose levels exhibited significant changes. The role of these candidate proteins is currently being validated (unpublished data). Thirdly, the research was conducted exclusively in vitro and in vivo using mouse models, and human data validation is lacking. Furthermore, the animal experiments utilized super-physiological doses of miR-293-5p mimics and pAd-TET1. To confirm these results and conclusions, future studies are required to include adipocyte-specific overexpression of TET1 in mice.

## Conclusion

Our study deciphers a hepatocyte-BAT regulatory axis where cold-induced exosomal miR-293-5p suppresses TET1 to activate thermogenesis, enhancing lipid oxidation and glucose disposal. Targeting hepatic miR-293-5p biogenesis offers novel therapeutic strategies against metabolic disorders.

## Materials and Methods

### Cell lines

Immortalized brown adipocytes, primary brown adipocytes, and beige adipocytes were cultured and differentiated according to established protocols [41,42,43]. Briefly, on day 0, confluent brown adipocytes were induced to differentiate using DMEM supplemented with 10% FBS, 20 nM insulin, 1 nM T3, 0.5 μM dexamethasone, 0.5 mM isobutylmethylxanthine, and 0.125 mM indomethacin for 48 hours. On day 2, the medium was changed to DMEM containing 10% FBS and 20 nM insulin, with 1 nM T3, for an additional 4 days to complete the differentiation process.

### Animal studies

All animal experiments were approved by the Binzhou Medical University Hospital Animal Care and Use Committee. Male C57BL6 mice, aged 6-8 weeks, were purchased from Jinan Pengyue Laboratory Animal Breeding Co., Ltd., Jinan, China. The mice were housed under a 12-hour light and 12-hour dark cycle at 22°C with ad libitum access to water and a standard rodent chow diet, unless otherwise specified.

An HFD, consisting of 60% of calories from fat, was obtained from Research Diets. HFD feeding commenced when the mice were 6 weeks old. Acute cold exposure was applied after 16 weeks of HFD feeding. To assess glucose tolerance and insulin sensitivity, mice underwent glucose tolerance tests (GTT) and insulin tolerance tests (ITT) following HFD. Mice were fasted for 16 hours before GTT or 5 hours before ITT, and then administered intraperitoneal injections of glucose (2 g/kg body mass) or insulin (0.75U kg^-1^ body mass), respectively. For indirect calorimetry, mice were acclimated in the Columbus Instruments Comprehensive Lab Animal Monitoring System, which measured energy expenditure and respiration under basal conditions. A scrambled agomiR was used as a control. The sequence for miR-293-5p agomiR is ACUCAAACUGUGUGACAUUUUG. For overexpression TET1, mice were injected bilaterally into the BATs with pAd-TET1 or pAd-GFP at a dose of 2×10^9^ PUF/kg in 0.1mL saline once a week for 1 month.

### Isolation of primary hepatocytes

Mice underwent in situ hepatic perfusion initiated with calcium/magnesium-free HEPES-buffered solution A (10 mM HEPES, 1 mM glucose, 0.2 μM EGTA, 0.2% BSA, pH 7.4) via the inferior vena cava for 3-5 min until hepatic decolorization (beige parenchyma). Following this, the perfusate was switched to digestion buffer B (PBS supplemented with 1 mM CaCl₂, 1 mM MgCl₂, 0.5 mg/mL collagenase D, 30 mM HEPES, and 0.2% BSA). Perfusion was discontinued upon observable capsular disruption, after which the liver was rapidly excised into ice-cold buffer A. The digested hepatic tissue was mechanically dissociated in buffer A, filtered through a 100 μm nylon mesh, and subjected to low-speed centrifugation (60 × g, 4°C, 6 min) to pellet parenchymal cells. The pellet underwent two washes with enzyme-free buffer B, followed by density gradient purification using 36% (v/v) Percoll™ (100 × g, 4°C, 10 min). After aspirating the supernatant, the hepatocyte-enriched fraction was resuspended in buffer B and plated onto collagen-coated culture plates with Williams’ Medium E containing 10% heat-inactivated fetal bovine serum and 1× antibiotic-antimycotic cocktail. A medium replacement protocol was executed after 16 h of primary culture to remove non-adherent debris.

### Isolation and characterization of hepatocyte derived exosomes

Primary hepatocytes were maintained in exosome-depleted DMEM (supplemented with 10% FBS pre-cleared by 120,000 × g centrifugation for 16 h) for 6 h under standard culture conditions (37°C, 5% CO₂). Following this acclimatization phase, the medium was replaced with fresh exosome-free DMEM for a 36-h secretory period. Cellular supernatants (20 mL pooled from 5×10⁶ cells) were processed through differential ultracentrifugation: initial clarification at 3,000 × g for 15 min (4°C) to remove cellular debris, followed by filtration through 0.22 μm pore-size polyethersulfone membranes to eliminate microvesicles >220 nm. The filtrate underwent ultracentrifugation at 120,000 × g (4°C, 60 min) using a Type 70 Ti rotor , with the resulting exosome pellet washed in ice-cold PBS and re-centrifuged under identical parameters to remove soluble protein contaminant. Exosome integrity was verified by transmission electron microscopy following negative staining with 2% uranyl acetate, revealing characteristic cup-shaped morphology with diameters of 30- 150 nm. Exosome morphology was assessed using transmission electron microscopy at the Electron Microscopy Center of Binzhou Medical University.

### Isolation of plasma exosomes and exosomal miRNAs

Plasma specimens underwent initial clarification at 3,000 × g (4°C, 15 min) to eliminate cellular debris. Following rapid thaw at 37°C, 250 μL aliquots were transferred to pre-chilled 1.5 mL low-binding tubes. Coagulation factors were removed by incubating supernatants with thromboplastin D (DiaPharma Group, 1:100 dilution) at 37°C for 15 min, followed by a secondary clarification at 10,000 × g (RT, 5 min). Cleaned supernatants were processed for extracellular vesicle isolation using ExoQuick™ Exosome Precipitation Solution combined with RNase A to degrade free-floating RNA. The mixture was incubated overnight at 4°C with gentle inversion. Prior to precipitation, murine RNase inhibitor was added to protect vesicular RNA integrity. EVs were pelleted at 1,500 × g and washed twice with ice-cold PBS. Exosomal miRNA extraction was performed using the miRNeasy Micro Kit with on-column DNase I treatment, strictly adhering to MIQE guidelines. RNA integrity was verified via Bioanalyzer 2100 with RIN >8.0, and quantified using a Qubit 4.0 Fluorometer.

### ChIP-reChIP

Primary immunocomplexes were eluted in 100 μL elution buffer (50 mM Tris pH 8.0, 1 mM EDTA, 1% SDS, 50 mM NaHCO3) at 65°C for 10 min. Beads were treated with 40 μL 10 mM DTT at 37°C for 30 min to dissociate residual antibodies. Combined eluates were diluted 10-fold in ChIP dilution buffer (16.7 mM Tris-HCl pH 8.0, 167 mM NaCl, 1.2 mM EDTA, 1.1% Triton X-100, 0.01% SDS) and subjected to secondary IP using anti-HDAC1 antibody under identical conditions.

### ChIP-qPCR

Cells were cross-linked with 1% formaldehyde for 10 min at 22-25°C. Chromatin was fragmented using a Covaris S220 Ultrasonicator (peak power: 140 W, duty factor: 5%, cycles/burst: 200) to generate 200-500 bp DNA fragments, verified by 2% agarose gel electrophoresis. Cleared lysates were divided into input controls and immunoprecipitation (IP) samples. IP samples were incubated overnight at 4°C with 5 μg of anti-Ty1 epitope antibody, anti-HDAC1, or species-matched IgG control. Protein-DNA complexes were captured by adding 20 μL pre-washed Dynabeads™ Protein G per reaction, followed by 1 h rotation at 4°C. Beads were successively washed in low-salt RIPA buffer (20 mM Tris-HCl [pH 8.0], 1 mM EDTA, 1% Triton x-100, 0.1% SDS, 140 mM NaCl, 0.1% Na deoxycholate), high-salt RIPA buffer (20 mM Tris-HCl [pH 8.0], 1 mM EDTA, 1% Triton x-100, 0.1% SDS, 500 mM NaCl, 0.1% Na deoxycholate), LiCl buffer (250 mM LiCl, 0.5% NP40, 0.5% Na deoxycholate, 1 mM EDTA, 10 mM Tris-HCl [pH 8.0]) and TE buffer (10 mM Tris-HCl [pH 8.0] and 1 mM EDTA). Crosslink reversal was performed in elution buffer (50 mM Tris-HCl pH 8.0, 1 mM EDTA, 100 mM NaCl, 0.5% SDS) containing 0.5 mg/mL proteinase K (Roche #03115887001) at 65°C for 4 h. DNA was purified by phenol:chloroform:isoamyl alcohol (25:24:1 v/v) extraction, followed by ethanol precipitation (2.5×volumes 100% ethanol, 0.1×volume 3 M NaOAc, -20°C overnight).Real-time qPCR primers are listed in Table S2. All data were normalized to input.

### miRNA sequencing

Total RNA was extracted using TRIzol and stored in diethyl pyrocarbonate-treated water at -20°C. miRNA libraries were prepared according to the manufacturer’s instructions using the NEBNext Illumina Small RNA Library Preparation Kit, which generates unique indexed libraries for each sample. Sequencing was performed on the Illumina HiSeq platform. The quality of the sequencing data associated with the sequencing core was assessed by the bioinformatics team.

### RNA-Seq Analysis

Total RNA was isolated from samples using the RNeasy Mini Kit (Qiagen, #74104), incorporating on-column DNase I digestion to remove genomic DNA contamination. RNA integrity was verified (RIN >8.0) prior to library preparation. Strand-specific RNA-seq libraries were constructed using the BGI RNA Library Prep Kit following poly(A)+ RNA selection, and sequenced (150 bp paired-end) on the BGISEQ-500 platform (average depth: 40 million reads/sample). Raw reads were quality-filtered and aligned to the mm10 genome (STAR v2.7.10a) using stringent parameters ("--outFilterMultimapNmax 1 --outSAMtype BAM SortedByCoordinate"). Mitochondrial RNA reads were excluded during alignment to prevent quantification bias. Gene-level counts were generated (featureCounts v2.0.1) against GENCODE M25 annotations.Differential expression analysis (edgeR v3.38.1) employed TMM normalization with significance thresholds (FDR<0.05, |log2FC|>1). Co-expressed gene modules were identified through hierarchical clustering (Ward’s method; Pearson correlation distance). Functional enrichment of DEGs (FDR<0.01) was performed using Enrichr (GO Biological Processes 2023; adjusted p<0.05).

### miRNA quantitative PCR

The exosomal RNAs used were poly(A)-tailed with poly(A) polymerase, followed by reverse transcription using modified oligo(dT) primers and SMART MMLV reverse transcriptase. Small RNA assays with miRNA-specific primers were used to convert RNA from other sources into cDNA via stem-loop reverse transcription. Quantitative PCR was conducted using the Applied Biosystems 7500 system with SYBR Green mix and primers specific to each miRNA. The relative abundance of each miRNA was calculated after normalization to U6, and results were expressed as fold changes relative to the control group.The primer sequences are shown in Table S2.

### Mimics/inhibitors transfection

Commercially available miRNA mimics, inhibitors, and negative controls (RiboBio, Guangzhou, China) were used. Transfections were carried out using iMAX reagent (Invitrogen) with 50 nM miRNA mimics/inhibitors or negative controls, according to the manufacturer’s protocol. Brown preadipocytes and primary hepatic s were seeded in 12-well plates at an 80% confluence and transfected with 25 nM nonspecific RNA control or miRNA mimics/inhibitors. Cells were maintained in fresh medium 12 hours post-transfection and subsequently harvested for Western blot or quantitative RNA analysis as required.

### Luciferase reporter gene assays

Various *Tet1* luciferase reporter plasmids (pTK81-IgK, 100 ng per transfection) were transiently transfected into HEK293 cells along with a Renilla luciferase vector (pRL, 10 ng per transfection). Cells were plated in 24-well plates and cultured for less than 24 hours before transfection. Constructs of 100 ng pMIR REPORT containing wild-type or mutated miR-293-5p binding sequences were used. Luciferase activity was measured using the Dual-Luciferase Reporter Assay System (Promega) and normalized to Renilla luciferase activity. Each experiment was conducted in triplicate and independently repeated at least three times.

### Co-culture experiments

Co-culture experiments were performed using 12-well transwell plates with 0.4 μm pore-sized filters (Corning Costar, USA) for 24 hours. Primary hepatic s were seeded in the transwell inserts, while brown preadipocytes were seeded in the bottom chamber.

### Oxygen Consumption

Freshly isolated adipose tissue deposits (approximately 40 mg) were minced in 1 mL phosphate-buffered saline supplemented with 25 mM glucose, 1 mM pyruvate, and 2% BSA. Cultured adipocytes were trypsinized, collected by centrifugation, and resuspended in the same buffer. Oxygen consumption was measured using a Clark electrode (Oxygraph+ System, Hansatech), and data were normalized according to tissue weight or total protein content.

### In vivo adeno-associated serotype 9 virus injection into BAT

Recombinant adeno-associated serotype 9 virus with TBG promoter for YBX1 knockdown ( AAV9-shYbx1) was purchased from Hanbio Biotechnology Co (Shanghai, China). The virus was diluted with sterile 1× PBS. We used AAV9-Ctrl as control. Five microliters of either AAV9-shYbx1 or AAV9-Ctrl was delivered into the BAT (total virus: 2×10^11^VG, injection volume: 200 µL, administered once every four weeks).

### In vivo adenovirus injection into BAT

Adenoviruses were purified by cesium chloride ultracentrifugation, and virus titers were determined in HEK293T cells by counting GFP-positive cells. Two-month-old male mice were used for viral injection. 100 µL of each adenovirus (5 × 10^9^ pfu) diluted in phosphate-buffered saline was injected bilaterally into the BAT. Mice were injected every 3 weeks for a total of two injections.

### Adenoviral expression vectors

The adenoviral expression vector pAd/CMV/V5-DEST, encoding TET1, was constructed following the manufacturer’s protocol. The TET1 sequence was amplified using the following primers: 5’- TATGTCGACATGTCTCGGTCCCGCCCCGCAAA-3’ (forward) and 5’- TAATTCTAGATTAGACCCAACGATTGTAGGGT-3’ (reverse).

### Serum triglyceride, total cholesterol and free fatty acids levels

Serum triglyceride, total cholesterol, and free fatty acids levels were assessed using a triglyceride assay kit according to the manufacturer’s instructions.

### Oil Red O staining

Mature adipocytes were washed once with phosphate-buffered saline and fixed in 4% buffered formaldehyde at room temperature. The cells were then stained with Oil Red O working solution for 1 hour. After staining, cells were washed several times with Milli-Q water and prepared for microscopic imaging.

### RNA preparation and quantitative real-time PCR

Total RNA from cells and tissues was isolated using TRIzol (Invitrogen). Reverse transcription was performed using HiScript® Q RT SuperMix (Vazyme) according to the manufacturer’s protocols. Quantitative real-time PCR (qPCR) was conducted using the ViiA 7 Real-Time PCR System (Applied Biosystems). The primer sequences are shown in Table S2.

### Western blot analysis

Cell or tissue samples were homogenized in cell lysis buffer [150 mM NaCl, 0.5% Triton-X-100, 5% glycerol, 50 mM Tris-HCl (pH 7.5), 1 mM PMSF, and a protease inhibitors mixture (Roche)]. Lysates were centrifuged at 13,000 g for 10 minutes at 4°C. Supernatants were quantified for protein concentration and analyzed by Western blotting with the indicated antibodies. The antibody information is listed in Table S3.

### Hematoxylin-Eosin Staining and Immunohistochemistry

Fresh tissue samples were fixed in 4% paraformaldehyde, embedded in paraffin, and stained with hematoxylin-eosin at room temperature for 24 hours. Histological images were captured using a microscope. For immunohistochemistry, slides were blocked with 2% horse serum, incubated overnight at 4°C with anti-UCP1 (1:150), and then washed. Slides were incubated with an appropriate peroxidase polymer-linked secondary antibody for 30 minutes, stained with 3,3’-diaminobenzidine substrate, and counterstained with hematoxylin.

### Data analysis

Data are presented as mean ± standard error of the mean (SEM). Differences between two groups were assessed using an unpaired two-tailed Student’s t-test. One-way analysis of variance (ANOVA) followed by Tukey’s test was used to evaluate the statistical significanceof three or more groups. Statistical significance is indicated as *p < 0.05, **p < 0.01, or ***p < 0.001.

### Data availability

All data, methods, and results of statistical analyses are reported in this paper and the associated Supplementary Materials. Any specific inquiries can be addressed to the corresponding author.

### Ethics and consent to participate

All animal experiments were approved and complied with the guidelines of the Institutional Animal Care and Use Committee of the Binzhou Medical University Hospital, China (permit number: 2022-G001-01).

### Consent for publication

Not applicable.

### Availability of data and materials

Not applicable.

### Competing interests

We declare there is no conflict interest for this manuscript.

## Funding

This work was supported by the National Natural Science Foundation of China (grant number: 82200981); the Natural Science Foundation of Shandong Province (grant numbers: ZR2022QH358, ZR202111250163); the Binzhou Medical University research start-up fund (grant number BY2021KYQD29); the Basic Research Project of Yantai Science and Technology Innovation and Development Plan (grant number: 2022JCYJ026); and the Special Funds of Taishan Scholars Project of Shandong Province (grant number: tsqn 202312384).

## Acknowledgements

We would like to thank Dr Dongning Pan and Ting Shi, Fudan University, for critical suggestions on the paper. We also thank Home for Researchers editorial team (www.home-for-researchers.com) for language editing service.

---

**Fig. S1.**
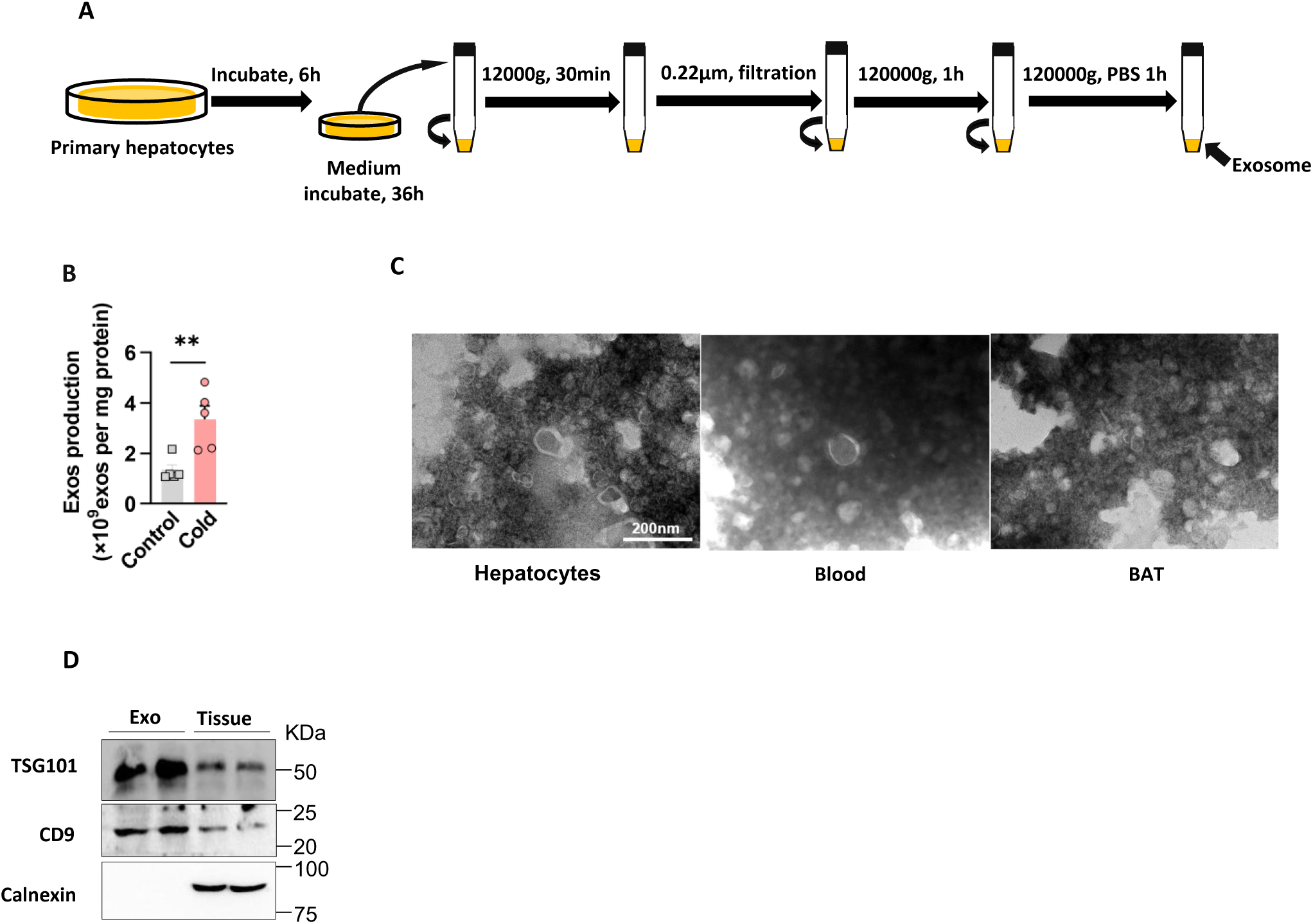
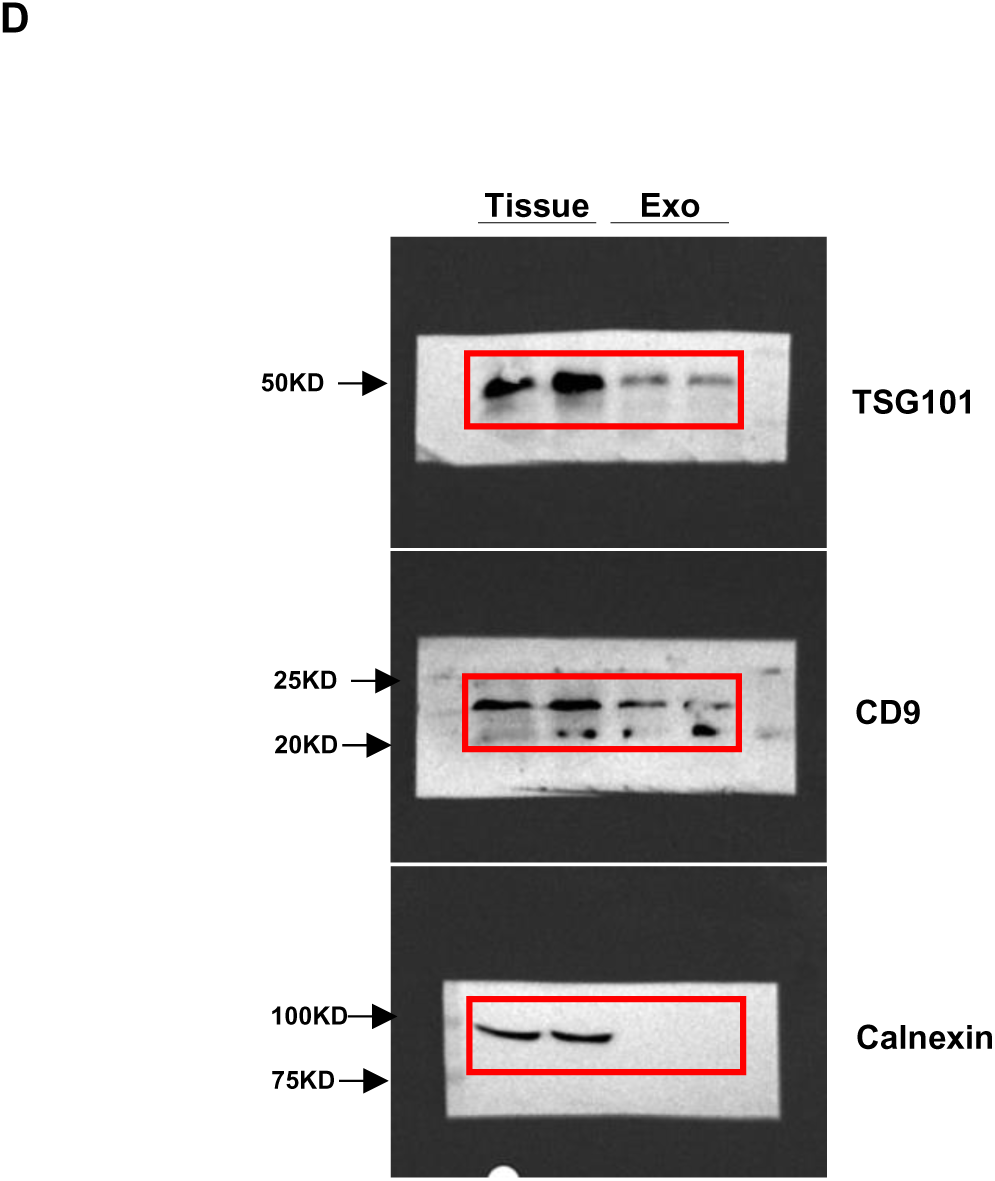
Stable of hepatocyte derived exosome after cold exposure. A. Flowchart illustrated the experimental procedures for exosome isolation from primary hepatocytes. B. Number of exosome from equal volumes of culture medium obtained primary hepatocytes from liver tissue. C. Representative electron micrographs of exosomes isolated from culture medium of hepatocytes, blood and BAT. Scale bar, 200 nm. D. Western blot analysis of CD9, TSG101 and Calnexin from equal volumes of culture medium obtained from primary hepatocytes and liver tissue.

**Fig. S2.**
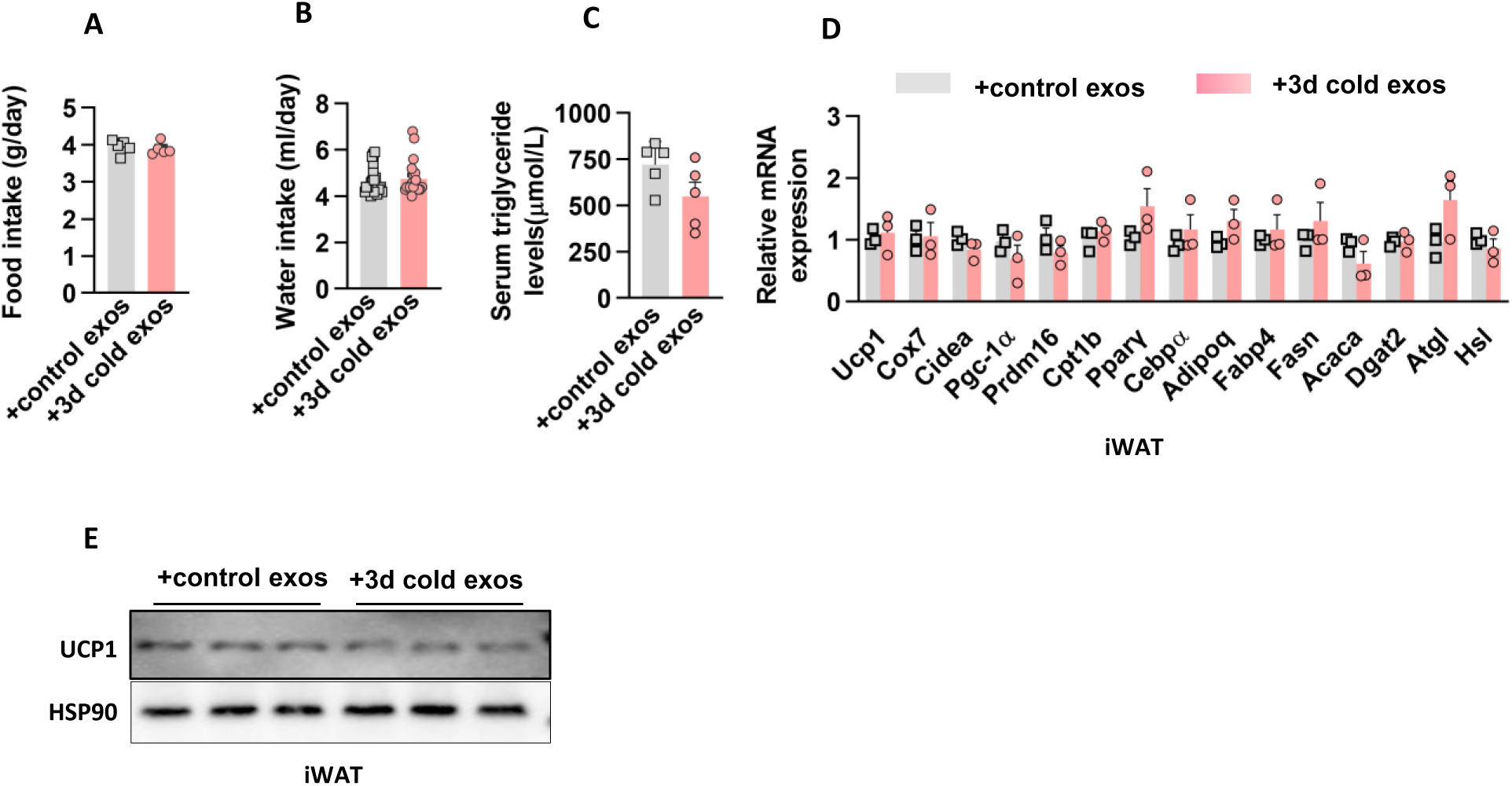
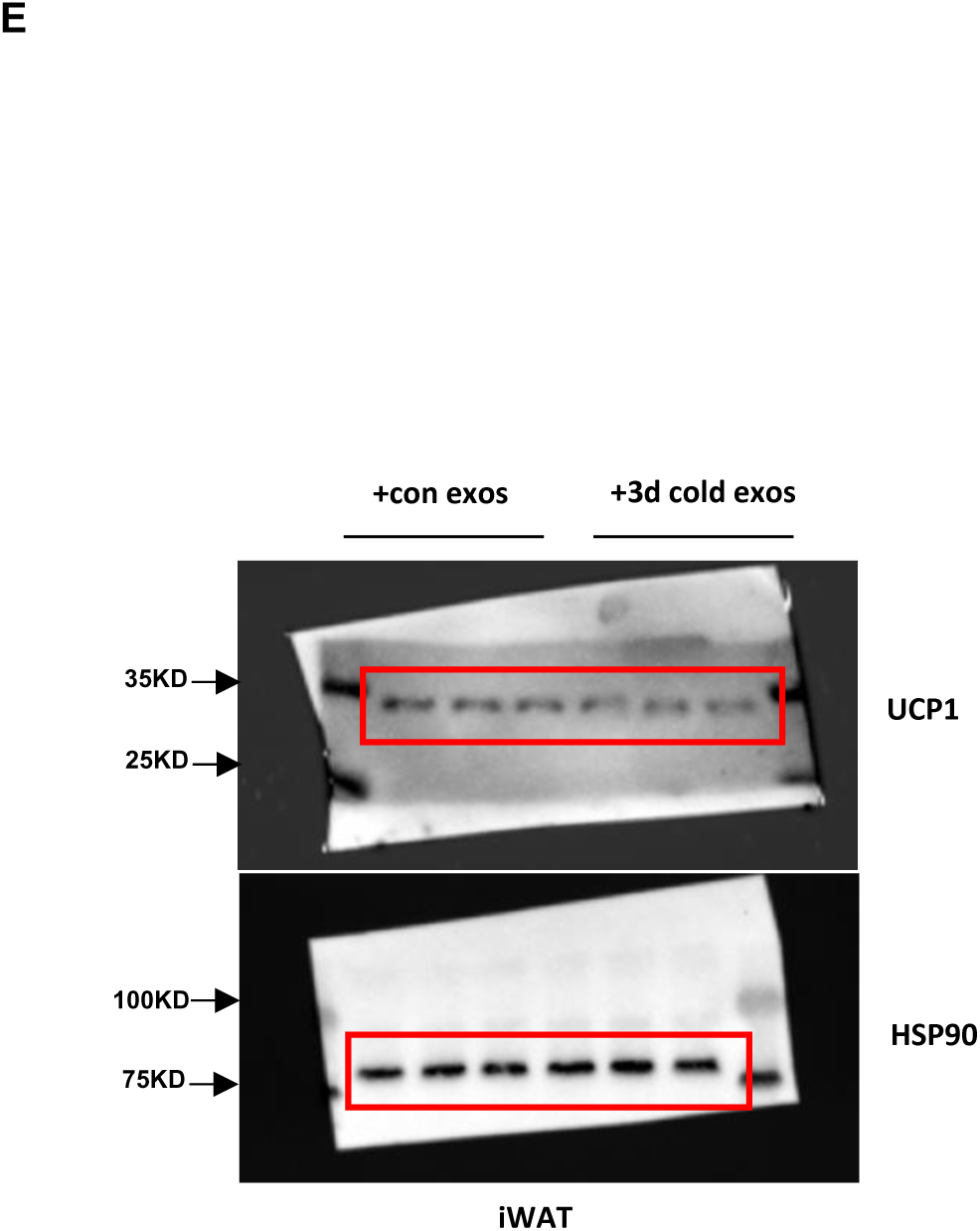
Hepatic-derived exosomes from cold-exposed mice did not affect the metabolism of iWAT. A. C57BL/6 male mice were used tail vein injection cold-exposed exosomes for 6 weeks. They consumed similar amount of water as control mice. n=5 mice each group. B. Chow diet food intake in mice of (A). n=5 mice each group. C. Serum triglyceride levels were assayed in mice of (A). n=5 mice each group. D. Gene expression in inguinal WAT from mice in (A). n=5 mice each group. E. UCP1 protein levels in iWAT from mice in (A).

**Fig. S3.**
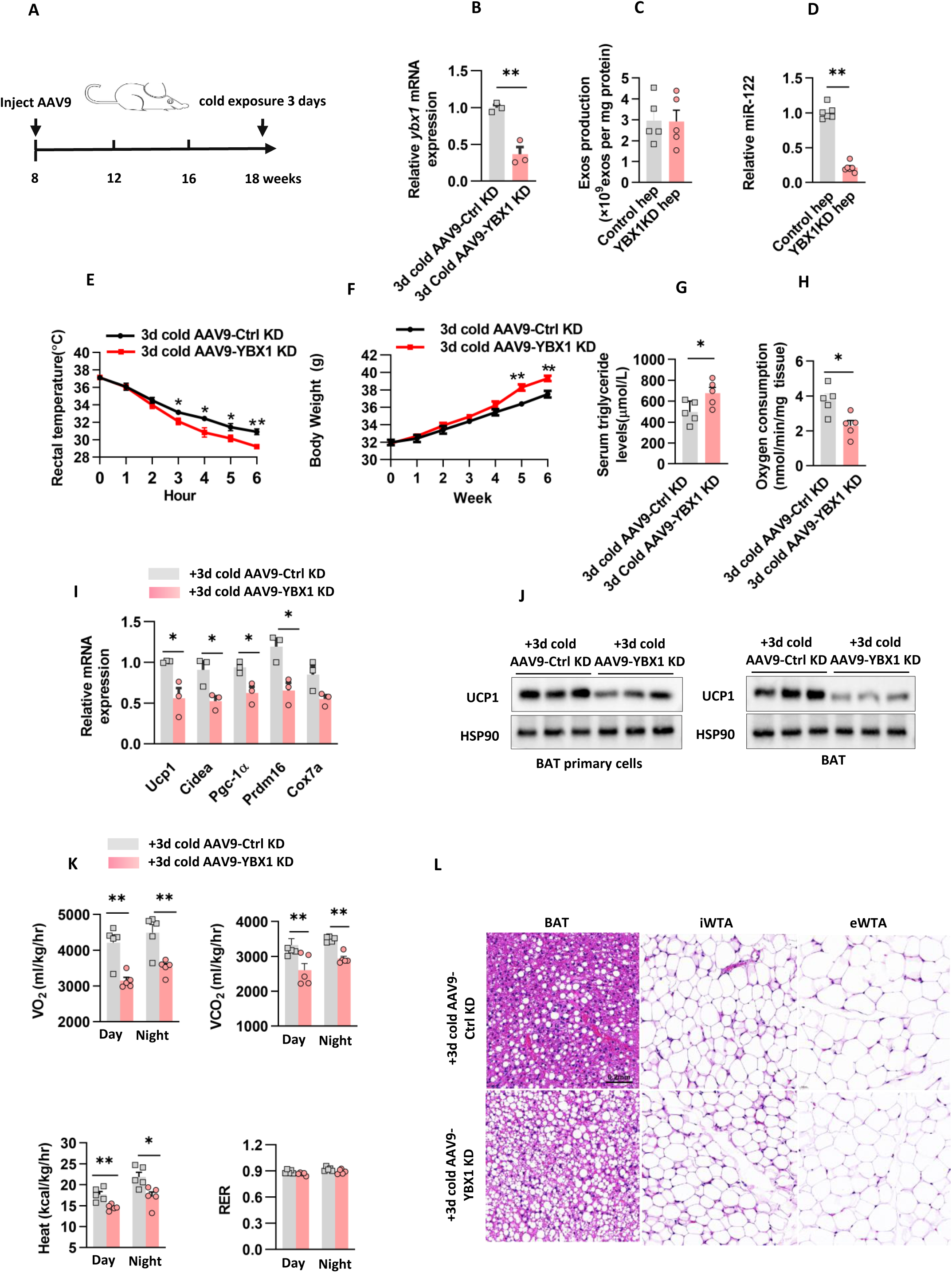
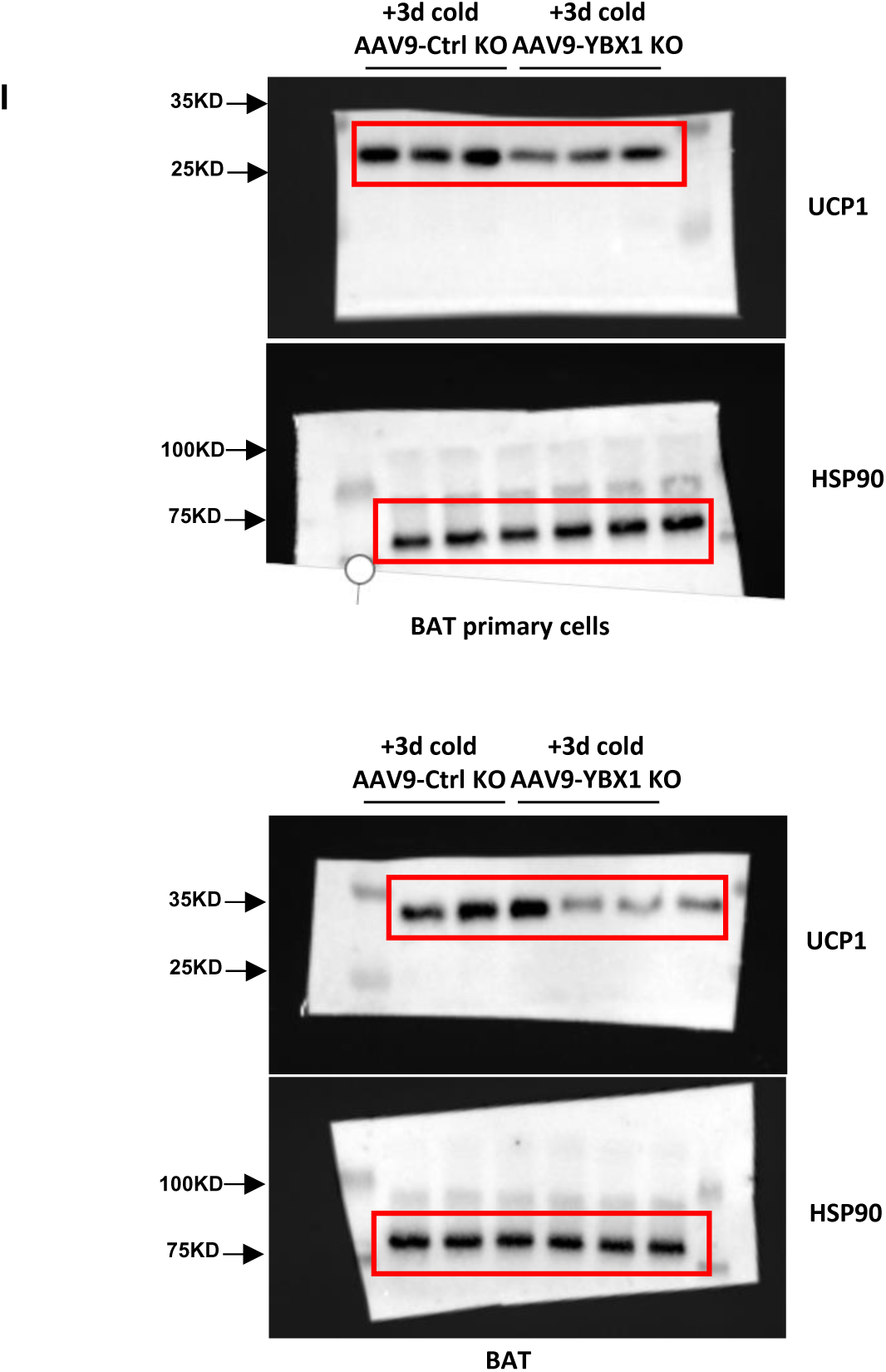
Decreased hepatocyte derived exosomes from cold-exposed mice was associated with thermogenic program BAT. A. The in vivo protocol that used tail vein injection AAV9-Ctrl KD or AAV9-YBX1 KD. B. Ybx1 gene expression analyzed by quantitative PCR in BAT as in (A). n=3 independent experiments. C. Effect of AAV9-YBX1 KD on hepatocyte derived exosomes production. D. miR-122 abundance within hepatocyte derived exosomes with and without AAV9-YBX1 KD. E. Rectal temperature was measured when mice in (A) were challenged to 4°C. n=5 mice each group. F. Body weight of mice in (A). n=5 mice each group. G. Serum triglyceride levels were assayed in mice of (A). n=5 mice each group. H. Oxygen consumption rate measured by Clark electrodes for BAT as in (A). n=5 independent experiments. I. Gene expression analyzed by quantitative PCR in BAT as in (A). n=5 independent experiments. J. UCP1 protein levels were analyzed in BAT primary cells and BAT. n=3 independent experiments. K. C57BL/6J male mice were subjected to the indirect calorimetry analysis at basal state. n=5 mice each group. L. H&E staining in (A) of mice from different treatment groups. Scale bar: 0.2mm for BAT,iWAT and eWAT. Data represent mean±SEM. *P<0.05; **P<0.01; ***P<0.001; unpaired two-tailed student t-test.

**Fig. S4.**
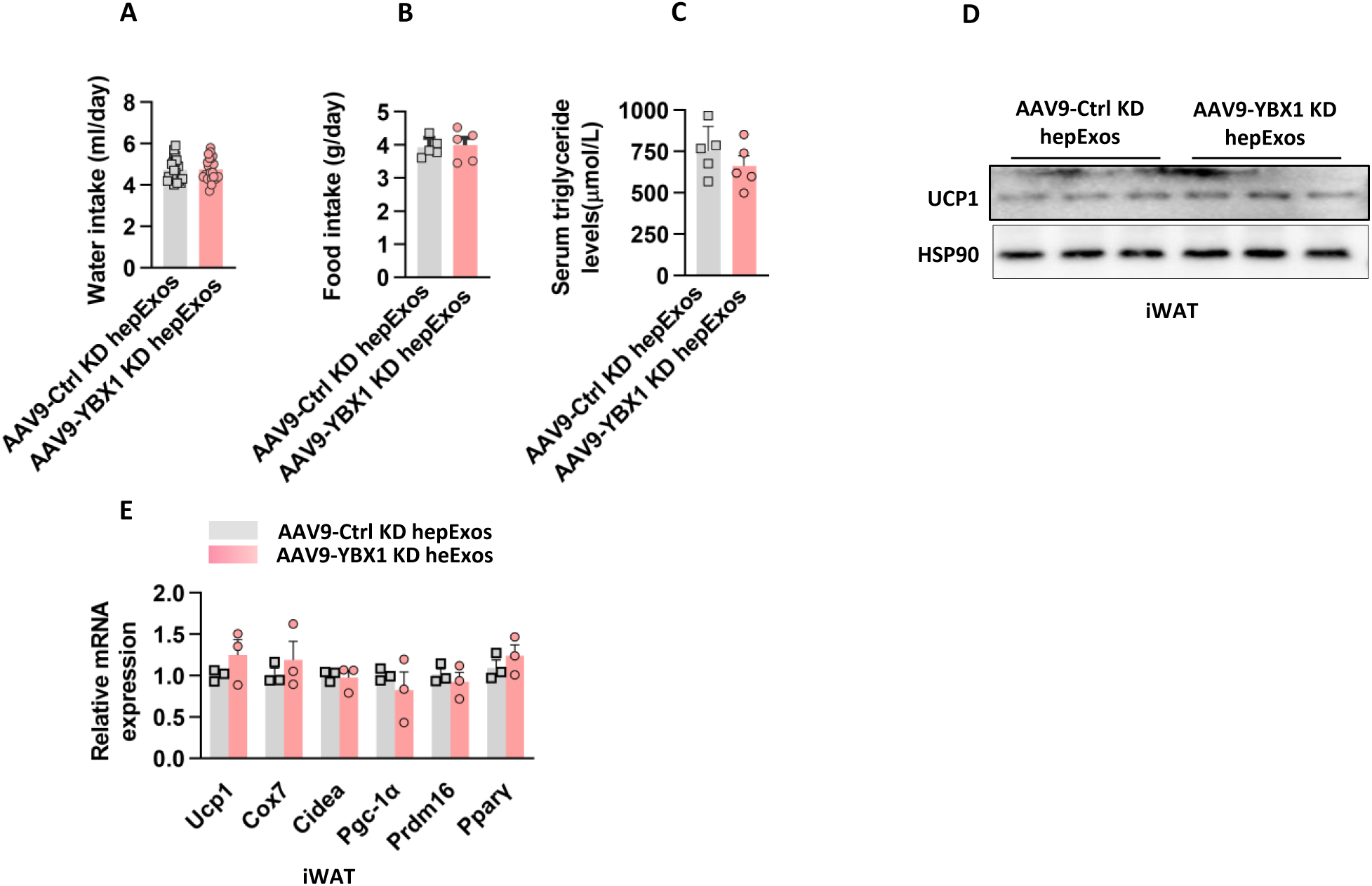
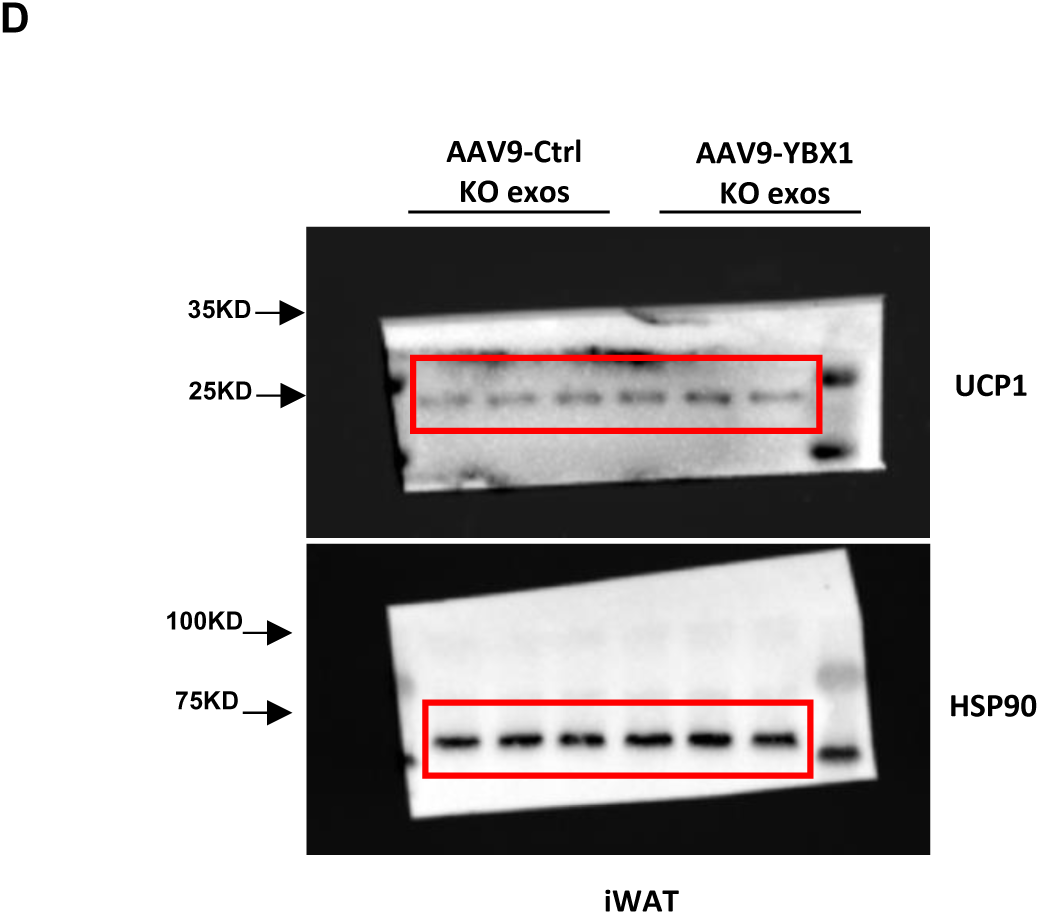
Hepatocyte derived exosomal miRNA acts on BAT to affect thermogenesis in vivo. A. C57BL/6 male mice were used intravenously injection AAV9-Ctrl KD or AAV9-YBX1 KD HepExos for 6 weeks. They consumed similar amount of water as control mice. n=5 mice each group. B. Chow diet food intake in mice of (A). n=5 mice each group. C. Serum triglyceride levels were assayed in mice of (A). n=5 mice each group. D. UCP1 protein levels in iWAT from mice in (A). E. Gene expression in inguinal WAT from mice in (A). n=5 mice each group.

**Fig. S5.**
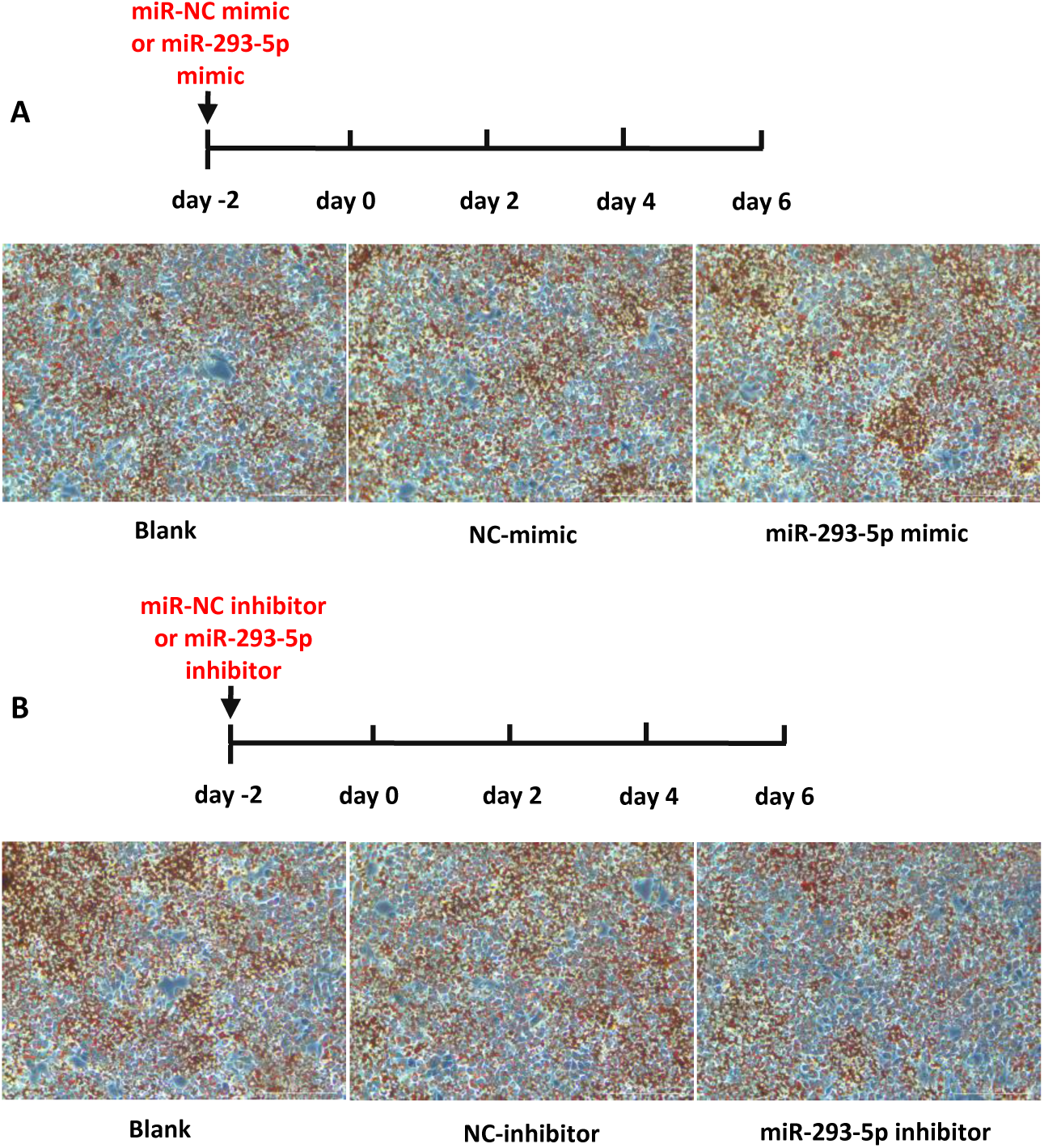
miR-293-5p without changing the differentiation of adipocyte. A. Immortalized brown adipocytes were transfected at day-2 with 25nM miR-293b-5p mimics or non-targeting control. Oil Red O staining of adipocytes differentiated from immortalized brown adipocytes on day 2. B. Immortalized brown adipocytes were transfected at day-2 with 25nM miR-293b-5p inhibitors or non-targeting control. Oil Red O staining of adipocytes differentiated from immortalized brown adipocytes on day 2.

**Fig. S6.**
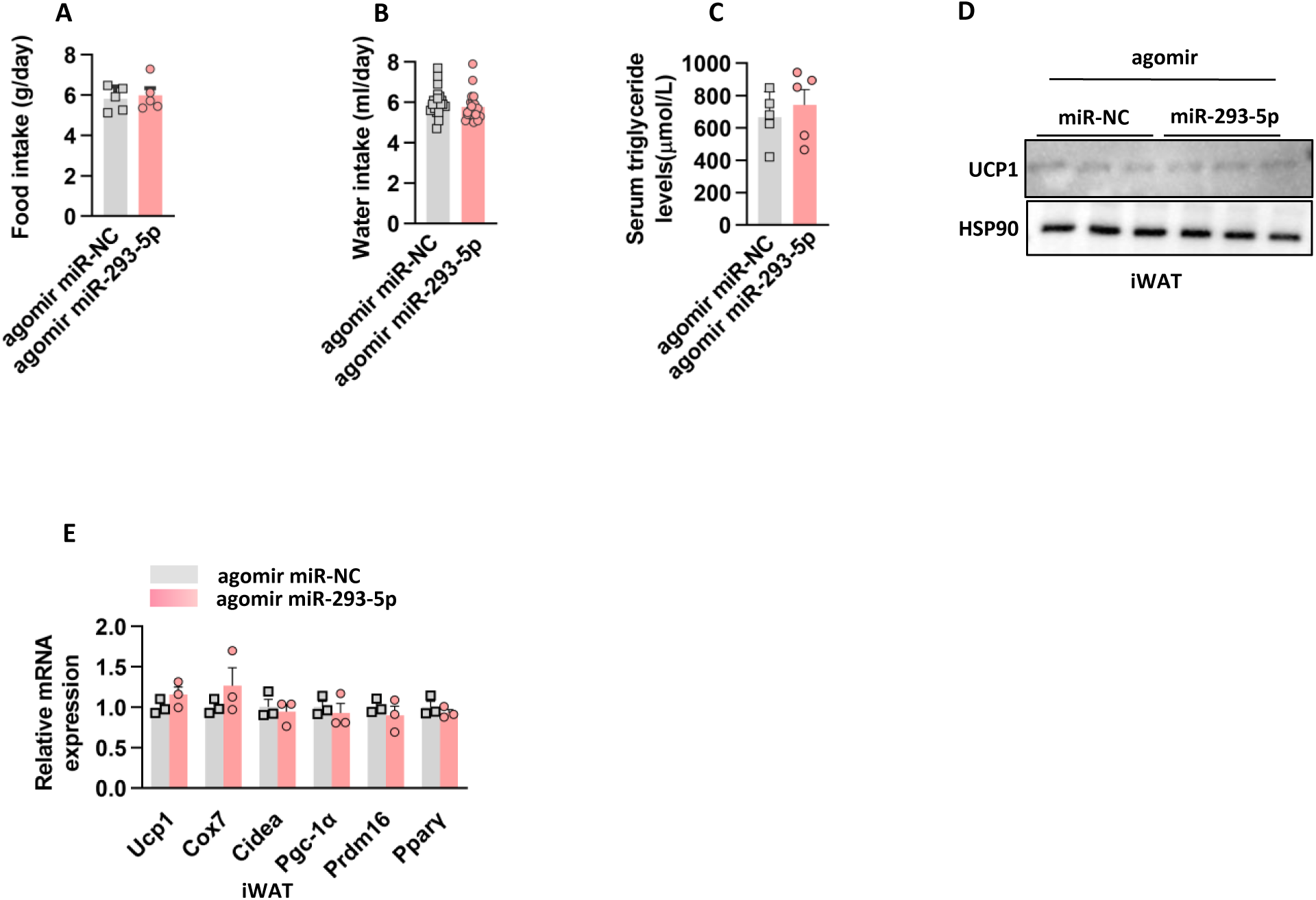
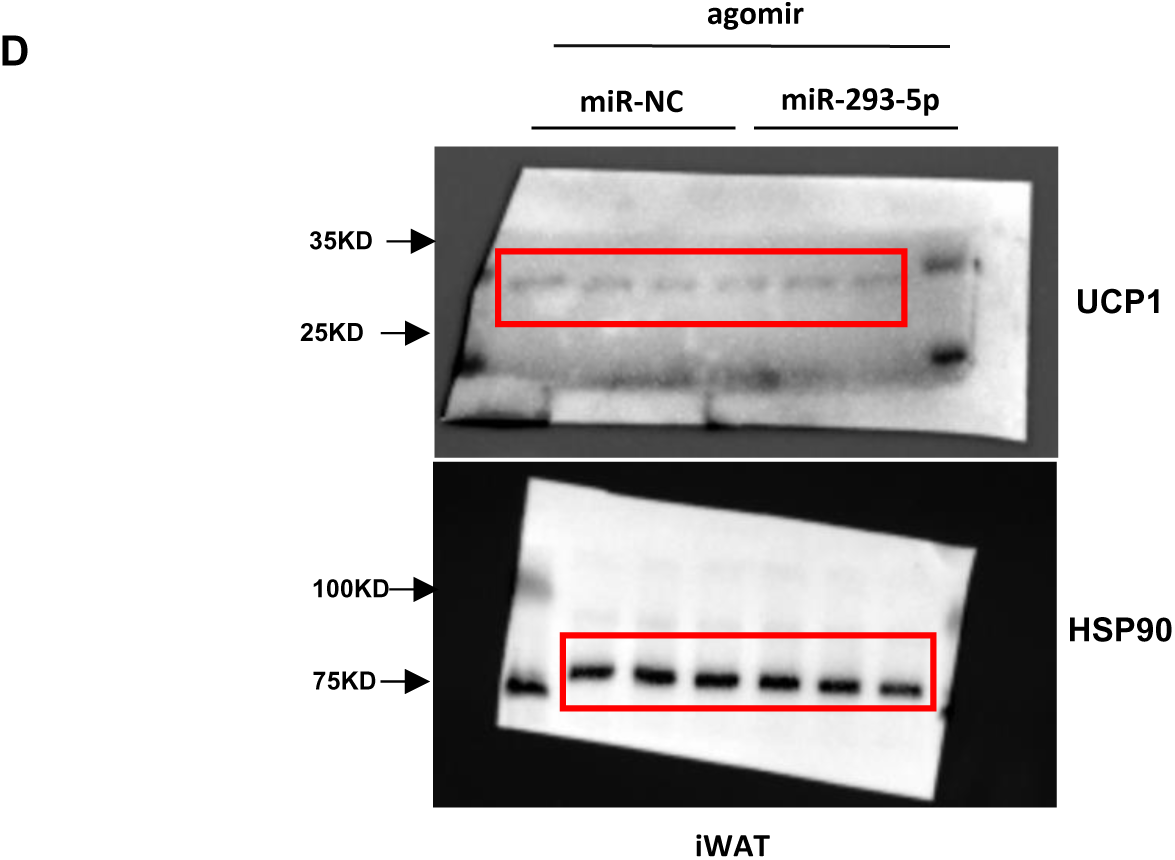
miR-293-5p injecting without affecting iWAT metabolism in mice. A. The consumed similar amount of water as control mice. n=5 mice each group. B. Chow diet food intake in mice of (A). n=5 mice each group. C. Serum triglyceride levels were assayed in mice of (A). n=5 mice each group. D. UCP1 protein levels in iWAT from mice in (A). E. Gene expression in iWAT from mice in (A). n=5 mice each group.

**Fig. S7.**
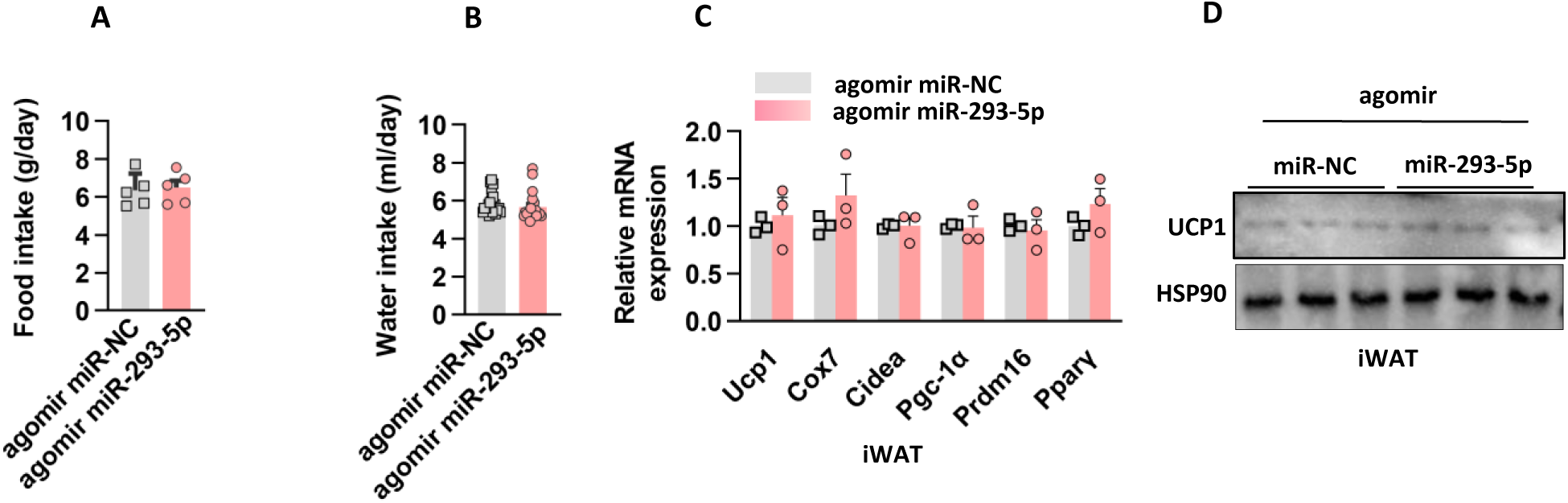
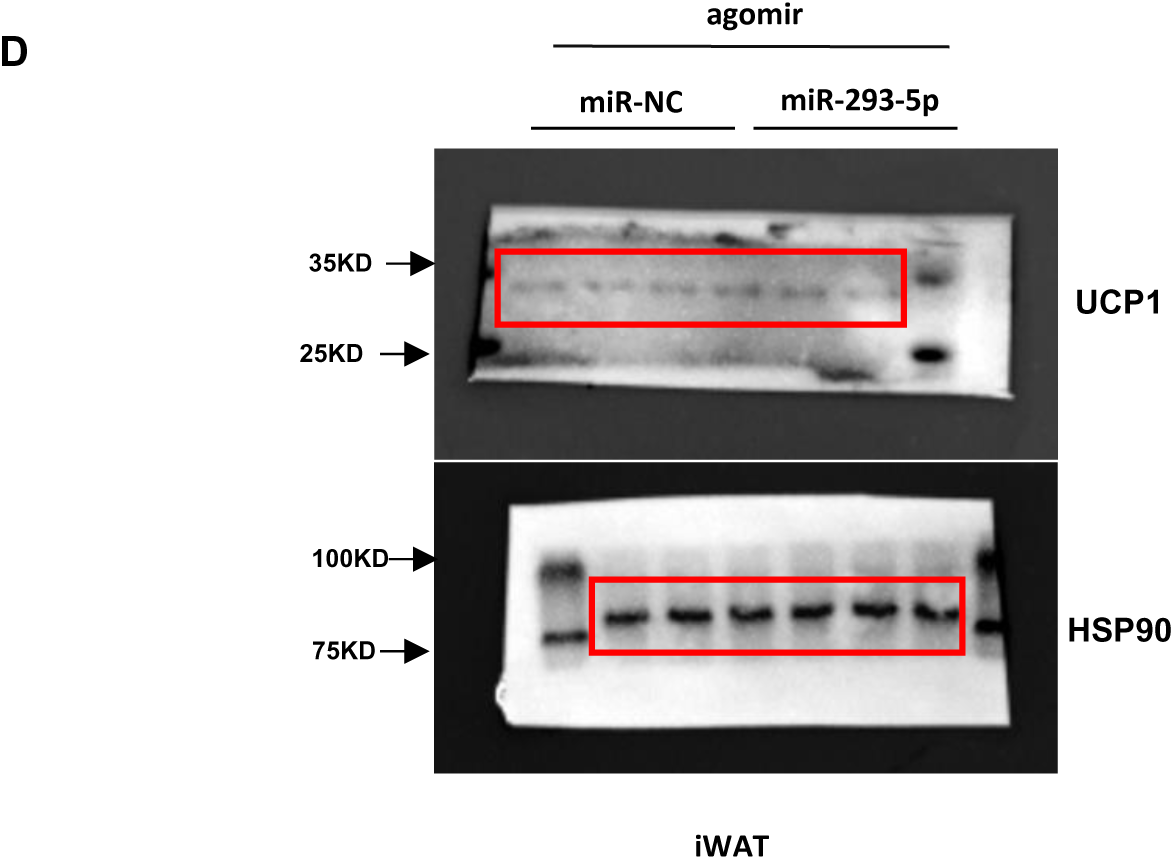
miR-293-5p injecting without affecting iWAT metabolism in diet-induced obesity mice. A. C57BL/6 male mice were used tail vein injection agomir miR-239-5p/miR-NC for 10 weeks. They consumed similar amount of water as control mice. n=5 mice each group. B. Chow diet food intake in mice of (A). n=5 mice each group. C. Gene expression in iWAT from mice in (A). n=5 mice each group.. D. UCP1 protein levels in iWAT from mice in (A).

**Table S2.**
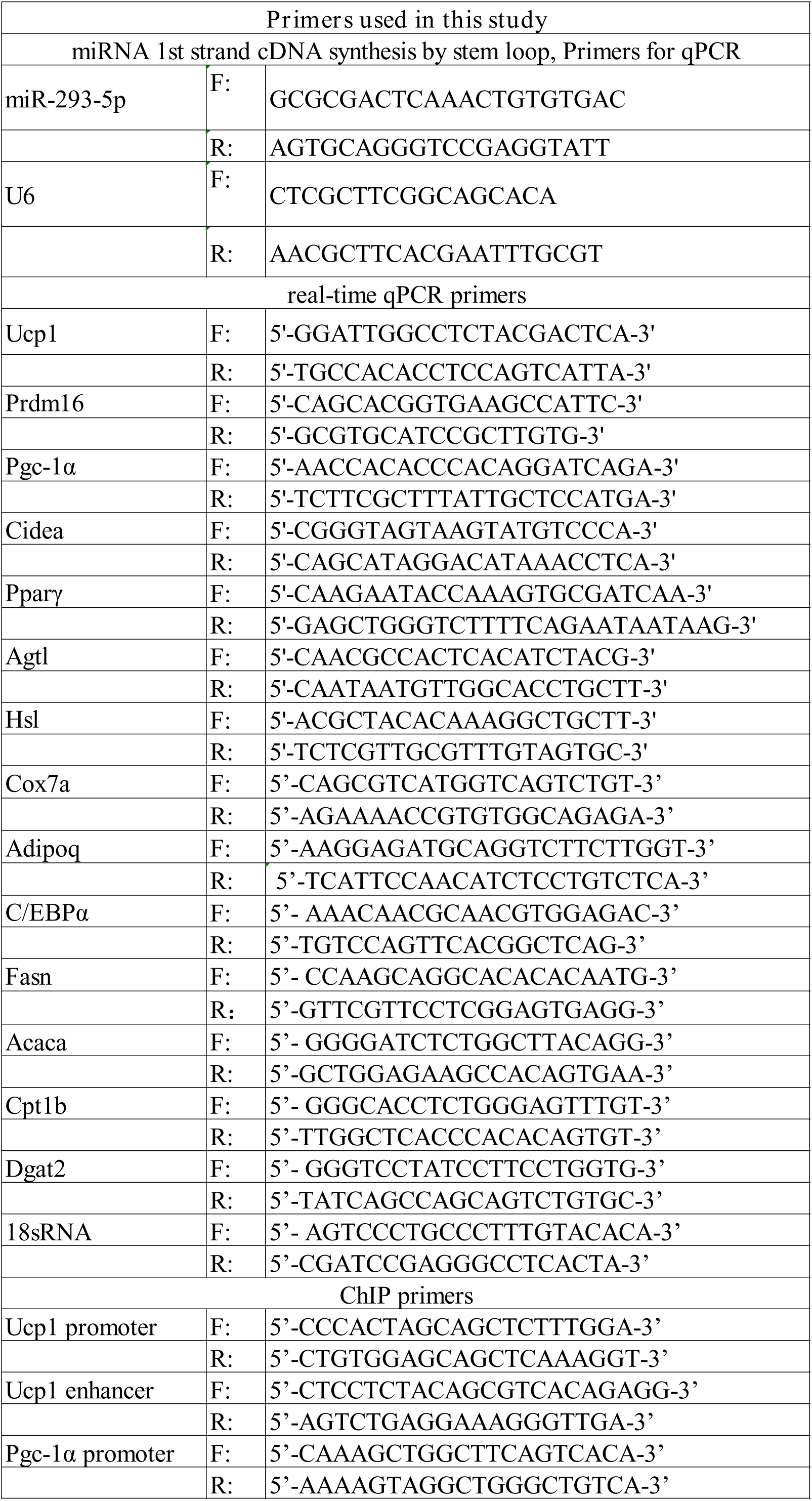

**Table S3.**

|  |  |  |
| --- | --- | --- |
| Company | Antibody | Product code |
| Abcam | Rabbit polyclonal anti-Ucp1 | Cat#ab10983 |
|  | Rabbit polyclonal anti-Tet1 | Cat#ab191698 |
|  | Rabbit polyclonal anti-Tsg101 | Cat#ab133586 |
|  | Rabbit polyclonal anti-CD9 | Cat#ab307085 |
|  | Rabbit polyclonal anti-Calnexin | Cat#ab22595 |
|  | Rabbit polyclonal anti-HDAC1 | Cat#ab280198 |
| Alpha Diagnostic International | Rabbit polyclonal anti-Ucp1 | Cat#UCP11-A |
| Cell Signaling Technology | Rabbit polyclonal anti-Pparγ1/2 | Cat#2443S |
| Proteintech | Rabbit polyclonal anti-α-Tubulin | Cat#66031-1-Ig |
|  | anti-Hsp90 | Cat#13171-1-AP |
| Company |  | Product code |
| Thermo Scientific | Thromboplastin D | 100356 |
| SBI | ExoQuick Exosome Precipitation Solution | 100356EXOQ20A-1 |
| Sigma-Aldrich | RNase A | R6513-10MG |
| NEB | Murine RNase inhibitor | M0314L |
| QIAGEN | miRNeasy Micro Kit | 217084 |
| Life Technologies | TRIzol | 15596026CN |
| BEYOTIME | Amplex Red triglyceride test kit | S0219S |
| BEYOTIME | Amplex Red free fatty acid test kit | S0215S |
| BEYOTIME | Amplex Red cholesterol and cholesterol ester test | S0211S |
| HyClone | Fetal bovine serum | SH30393.03 |
| Vazyme | ChamQ Universal SYBR Qpcr Master Mix | Q711 |
| Vazyme | RNA isolater Total RNA Extraction Reagent | R401 |
| Sigma-Aldrich | Insulin | I3536 |
| Sigma-Aldrich | T3 | T2877 |
| Sigma-Aldrich | IBMX | I5859 |
| Sigma-Aldrich | Indomethacin | I7378 |
| Sigma-Aldrich | Dexamethasone | D4902 |

| **BAT** | Cold1_count | Cold2_count | Cold3_count | Cold4_count | Control1_count | Control2_count | Control3_count | Control4_count |
| --- | --- | --- | --- | --- | --- | --- | --- | --- |
| mmu-miR-5106 | 230 | 218 | 360 | 446 | 66 | 176 | 126 | 342 |
| mmu-miR-293-5p | 1076 | 1680 | 1944 | 1734 | 242 | 395 | 397 | 345 |
| mmu-miR-3960 | 236 | 264 | 9149 | 7010 | 321 | 342 | 277 | 235 |
| mmu-miR-6374 | 20 | 16 | 28 | 32 | 5 | 17 | 19 | 16 |
| mmu-let-7f-5p | 405 | 316 | 579 | 487 | 1309 | 3094 | 1414 | 2456 |
| mmu-let-7a-5p | 662 | 462 | 1111 | 801 | 2540 | 5818 | 2616 | 2314 |
| mmu-let-7e-5p | 63 | 61 | 138 | 106 | 283 | 639 | 379 | 368 |
| mmu-miR-2137 | 194 | 229 | 456 | 489 | 1164 | 1155 | 1851 | 1765 |
| mmu-let-7d-5p | 161 | 133 | 225 | 142 | 535 | 895 | 638 | 765 |
| mmu-let-7c-5p | 2266 | 2457 | 3506 | 3085 | 7452 | 14675 | 10427 | 23143 |
| mmu-let-7g-5p | 180 | 144 | 240 | 190 | 534 | 928 | 556 | 657 |
| mmu-miR-98-5p | 10 | 7 | 24 | 20 | 47 | 74 | 62 | 73 |

| **BLOOD** | Cold1_count | Cold2_count | Cold3_count | Cold4_count | Control1_count | Control2_count | Control3_count | Control4_count |
| --- | --- | --- | --- | --- | --- | --- | --- | --- |
| mmu-miR-5126 | 234 | 234 | 117 | 94 | 817 | 594 | 632 | 359 |
| mmu-miR-6938-3p | 28 | 33 | 40 | 46 | 3 | 2 | 1 | 0 |
| mmu-miR-2137 | 120 | 136 | 124 | 120 | 523 | 476 | 294 | 235 |
| mmu-miR-3473b | 1680 | 1187 | 484 | 405 | 3024 | 2261 | 3511 | 1977 |
| mmu-miR-3473e | 1661 | 1168 | 476 | 398 | 2952 | 2225 | 3469 | 1934 |
| mmu-miR-5130 | 109 | 102 | 34 | 16 | 317 | 210 | 292 | 88 |
| mmu-miR-3106-5p | 68 | 65 | 96 | 107 | 32 | 16 | 10 | 8 |
| mmu-miR-5128 | 15 | 10 | 2 | 4 | 28 | 20 | 32 | 36 |
| mmu-miR-5106 | 642 | 652 | 104 | 166 | 83 | 69 | 80 | 52 |
| mmu-miR-16-5p | 1190 | 1185 | 1119 | 1333 | 98 | 79 | 46 | 26 |
| mmu-miR-12193-5p | 13 | 11 | 6 | 10 | 28 | 27 | 18 | 11 |
| mmu-miR-6990-5p | 137 | 100 | 111 | 128 | 58 | 32 | 48 | 36 |
| mmu-miR-29a-3p | 27 | 40 | 19 | 29 | 17 | 15 | 2 | 5 |
| mmu-miR-293-5p | 2211 | 2611 | 1623 | 1832 | 80 | 140 | 40 | 30 |

| **LIVER** | Cold1_count | Cold2_count | Cold3_count | Control1_count | Control2_count | Control3_count |
| --- | --- | --- | --- | --- | --- | --- |
| mmu-miR-122-5p | 42980 | 101065 | 55153 | 8746 | 13122 | 8889 |
| mmu-miR-16-5p | 102 | 247 | 179 | 24 | 16 | 53 |
| mmu-miR-293-5p | 12833 | 13232 | 15773 | 415 | 471 | 300 |
| mmu-miR-7665-5p | 35 | 63 | 33 | 164 | 151 | 124 |
| mmu-miR-674-5p | 14 | 22 | 12 | 7 | 0 | 4 |
| mmu-miR-7a-5p | 16 | 32 | 24 | 5 | 11 | 2 |
| mmu-miR-6769b-5p | 13 | 24 | 17 | 11 | 4 | 15 |
| mmu-miR-504-3p | 10 | 12 | 10 | 29 | 37 | 25 |
| mmu-miR-7081-5p | 31 | 72 | 46 | 15 | 31 | 52 |
| mmu-miR-7669-3p | 17 | 35 | 25 | 36 | 29 | 52 |

